# The SH3BGRL protein family nucleates and caps actin filaments via its conserved thioredoxin fold

**DOI:** 10.1101/2025.03.25.645244

**Authors:** Robin S. Heiringhoff, Daniel Marke, Ute Curth, Johannes N. Greve

## Abstract

Cellular actin polymerization is a tightly regulated process, typically controlled by proteins with specialized domains such as the Wiskott-Aldrich syndrome protein homology 2 (WH2) domain. Here, we identify SH3BGRL family proteins as modulators of actin dynamics, uniquely characterized by their thioredoxin (Trx) fold structure and the absence of the canonical CXXC enzymatic site essential for redox activity. The Trx fold is generally associated with enzymatic activity; however, in this context, it functions non-enzymatically to enhance actin filament nucleation and inhibit depolymerization. The family member SH3BGRL-2 was previously identified as part of the spectrin–actin complex in porcine erythrocytes. Further structural analysis reveals that human SH3BGRL proteins share structural homology with the C-terminal region of *Saccharomyces cerevisiae* YFR016c/Aip5, a known actin nucleation factor reported to bind G-actin. Notably, our results show that human SH3BGRL proteins do not bind G-actin directly. While they do not interact with G-actin, SH3BGRL proteins significantly increase actin assembly rates by accelerating filament nucleation without affecting barbed end elongation, as demonstrated in pyrene-actin bulk-polymerization assays and total internal reflection fluorescence microscopy (TIRFM) based single-filament studies. Furthermore, using all-atom molecular dynamics (MD) simulations and *in vitro* assays that directly probe the pointed end of the actin filament, we show that SH3BGRL proteins inhibit the depolymerization of existing filaments by interacting with the pointed end of the actin filament, also in the presence of the well-characterized pointed end capping protein tropomodulin. Our results indicate that all SH3BGRL family proteins promote actin nucleation by stabilizing energetically unstable actin dimers and trimers and inhibit depolymerization by direct association with the pointed end, suggesting a direct role for the Trx fold in actin dynamics.

## Introduction

The thioredoxin (Trx) protein fold is one of the most widely conserved structural motifs, present across all domains of life. Its characteristic three-dimensional structure consists of a central four-stranded β-sheet flanked by four α-helices [1,2]. The best-known protein families that contain one or more Trx fold domains include thioredoxins, protein disulfide-isomerases, glutaredoxins, bacterial thiol-disulfide oxidoreductases (Dsb), glutathione S-transferases, and glutathione peroxidases, which are involved in processes such as protein folding, protection against oxidative stress, and redox signaling, among others [1–3]. While the first four are redox enzymes that use the canonical dithiol CXXC motif in their active sites to reduce substrates, the latter two have a monothiol active site (CXXS/T). In addition to these well-studied protein families, many other Trx fold-containing proteins with uncharacterized molecular functions have been described in recent years [2].

Members of the SH3-binding glutamic acid rich (SH3BGR) protein family are small proteins, ranging from 10 kDa to 20 kDa in size. They are characterized by a common Trx fold, but notably lack a catalytic CXXC or CXXS/T motif canonically associated with Trx fold containing enzymes [1]. The eponymous member of the family, the cardiac and skeletal muscle-specific SH3BGR, is further distinguished by a C-terminal extension that is rich in glutamic acid residues [4,5]. The remaining three members of the SH3BGR family, named SH3-binding glutamic acid rich-like /-2 /-3 (SH3BGRL/-2/-3), are ubiquitously expressed and lack the C-terminal extension [6–8]. SH3BGR, SH3BGRL and SH3BGRL-2 all feature a single proline-rich sequence (PLPPQIF), which contains the eponymous SH3-binding motif (PXXP) and a Homer EVH1 binding motif (PPXXF). The shortest family member, SH3BGRL-3, contains only the Homer EVH1 binding motif [9,10]. In all cases binding of proteins to these motifs is unlikely, as the motifs are buried inside the tertiary structure [10] (**Figure 1B**).

**Figure 1:**
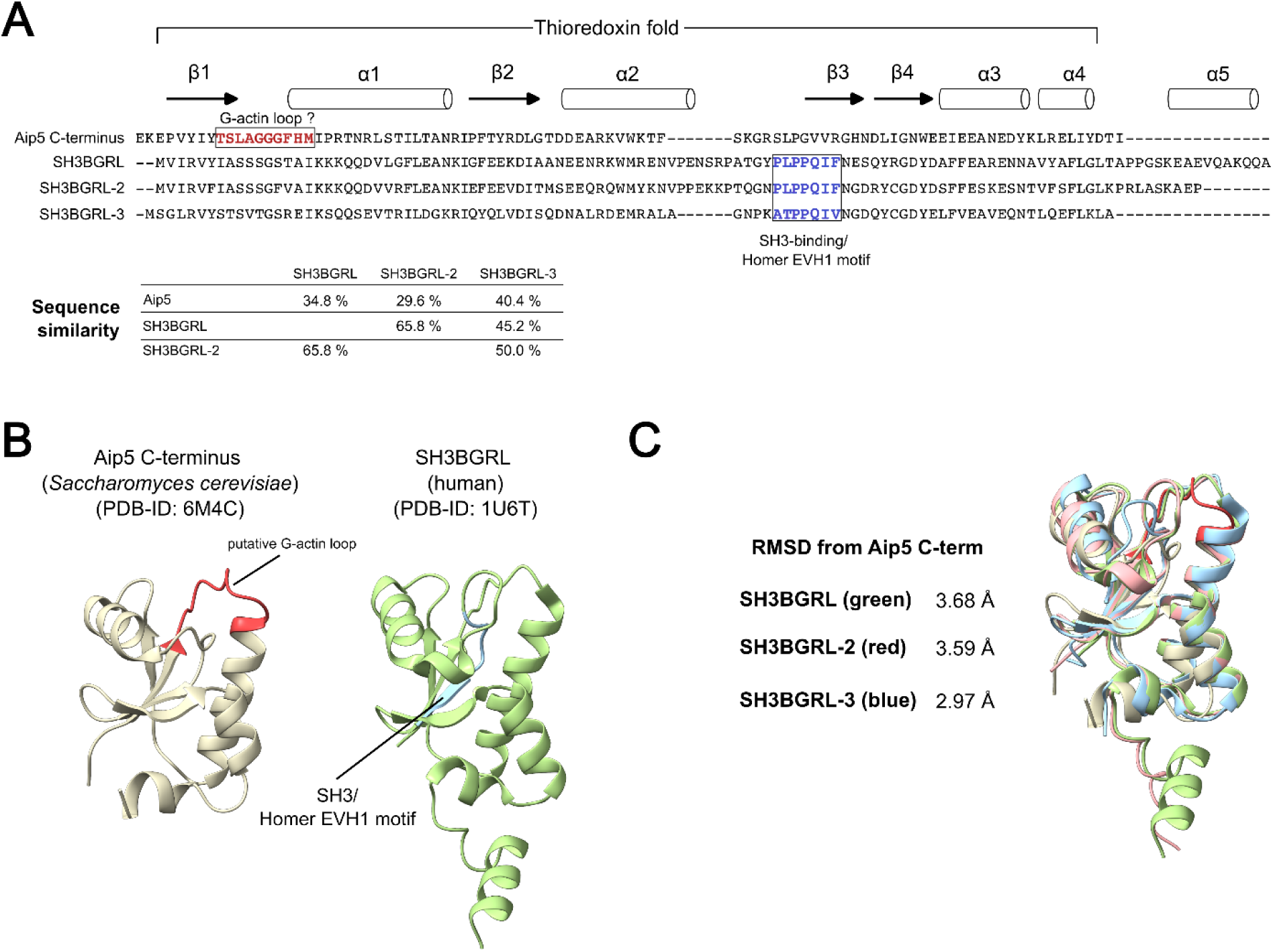
Sequence and structure comparison of the human SH3BRGL isoforms and the C-terminal region of Aip5 from *Saccharomyces cerevisiae*. **(A)** Sequence alignment of the human SH3BGRL isoforms and the Aip5 C-terminus (generated using Clustal Omega [61]). The putative G-actin loop in Aip5 and the SH3-binding/Homer EVH1 motif in SH3BGRL isoforms are highlighted in the sequence. Sequence similarities were determined using the Sequence Manipulation Suite [62] and given in the table. **(B)** Published experimental structures of the Aip5 C-terminus from *Saccharomyces cerevisiae* and of human SH3BGRL. The putative G-actin loop in Aip5 and the SH3-binding/Homer EVH1 motif in SH3BGRL are highlighted in the structure. **(C)** Structural alignment of the Aip5 C-terminus with the human SH3BGRL isoforms using the experimental structures of Aip5 C-terminus, SH3BGRL, SH3BGRL-3 and an AlphaFold-generated model of human SH3BGRL-2. The RMSD values were determined using the matchmaker function in ChimeraX [53].

The absence of the catalytic motif in the otherwise structurally conserved Trx fold raises the question of the functional relevance of the SH3BGR protein family. Previous studies on the family member SH3BGR indicated that production of the protein is important for proper sarcomere organization and maintenance [5,11]. Studies on the ubiquitously produced isoforms mainly place the function of the proteins in the context of carcinogenesis, either reporting expression as oncogenic [12–14] or anti-oncogenic [12,15,16], depending on isoform and species. Furthermore, studies in zebrafish indicate relevance of SH3BGRL for female fertility [17] and of SH3BGRL, SH3BGRL-2 and SH3BGRL-3 for organogenesis throughout different stages of development [18]. While members of the SH3BGR family are often described as adaptor proteins mediating protein-protein interactions due to the presence of a SH3 binding motif in their primary sequence, experimental data on the direct interaction between the protein family and binding partners are limited. A study reported direct interaction between SH3BGRL-3 and human myosin-1C by performing co-immunoprecipitation and mass spectrometry [19]. The same study reported that knocking down SH3BGRL-3 reduced the migratory capacity of MDA-MB-231 breast carcinoma cells, suggesting a potential link between SH3BGRL proteins and actin dynamics in non-muscle cells. In addition, direct interaction between SH3BGRL and ribosomal subunits is reported [14]. Cryo-electron microscopy combined with mass spectrometry has provided direct visual evidence of the binding of SH3BGRL-2 to actin filaments within the native spectrin–actin complex isolated from porcine erythrocytes [20]. In this multi-protein complex, SH3BGRL-2 is located at the pointed end of the actin filament alongside the capping protein tropomodulin, where it is thought to act as an additional capping protein to protect and stabilize the pointed end. This finding appears to contradict the results of previous fluorescence microscopic studies on the homologous SH3BGR isoform in *Xenopus laevis* heart muscle. These earlier studies showed SH3BGR localized to the Z-disc of the sarcomere, where the actin capping protein CapZ stabilizes the barbed ends of the filaments [5].

The widespread production of the SH3BGRL isoforms and the evident direct interaction of the SH3BGRL-2 isoform with actin filaments indicate a direct involvement of these proteins in the maintenance and regulation of cytoskeletal actin dynamics, extending beyond erythrocytes to other non-muscle cells. In this study, we utilize an integrated approach combining bioinformatics and *in vitro* biochemical analysis with purified proteins to explore the potential roles of human SH3BGRL isoforms in modulating actin dynamics.

## Results

### SH3BGRL proteins and the C-terminal region of actin-interacting protein 5 (Aip5) from *Saccharomyces cerevisiae* share a common thioredoxin fold

Actin-binding proteins utilize a variety of conserved protein folds to interact with both monomeric (G-) actin and filamentous (F-) actin. Among the most common folds are the actin-depolymerizing factor/cofilin (ADF/cofilin) domain fold, the Wiskott-Aldrich syndrome protein (WASP) homology-2 domain (WH2) fold, the gelsolin-homology domain fold, and the myosin motor domain fold [21]. The thioredoxin (Trx) fold, however, is not widely recognized as a major actin-binding fold.

We initially searched the BioGRID database to identify binding partners of the SH3BGRL protein family associated with the actin cytoskeleton [22]. The BioGRID database yields 150 unique interactors for SH3BGRL, 29 unique interactors for SH3BGRL-2 and 57 unique interactors for SH3BGRL-3. Closer inspection of the datasets reveals numerous proteins directly associated with the actin cytoskeleton in the datasets for SH3BGRL and SH3BGRL-2 e.g. the cytoskeletal actin isoforms β- and γ-actin, tropomodulin isoforms, subunits of the Arp2/3 complex and various cytoskeletal myosin isoforms (**Suppl. Table 1)**. None of these cytoskeleton-associated proteins are present in the available dataset for SH3BGRL-3. The available interactomes, which detail interactions between SH3BGRL family proteins and actin or actin-associated proteins, are based on primary data obtained through affinity capture-mass spectrometry from HCT116 and HKe-3 colon cancer cell lines, as well as HEK293 cells [23–25]. These findings suggest that the SH3BGRL protein family interacts with the actin cytoskeleton in cells that do not possess the specialized spectrin–actin complex found in erythrocytes. [20].

Next, to identify potential structural homologues implicated in actin dynamics, we conducted a protein structure search in the Protein Data Bank (PDB) using the SH3BGRL-2 sequence as the query in Foldseek [26]. Among many thioredoxin and glutaredoxin protein structures, we identified the ordered C-terminal region of actin-interacting protein 5 (Aip5) from *Saccharomyces cerevisiae* (PDB ID: 6M4C) as a structural homologue to SH3BGRL-2, with a template modeling score of 0.77, an RMSD of 2.63 Å, and 21.2% sequence identity (**Figure 1B**).

A structural alignment of the available crystal structure of the Aip5 C-terminus with the crystal structures of human SH3BGRL (PDB ID: 1U6T), human SH3BGRL-3 (PDB ID: 1T1V), and an AlphaFold-generated model of human SH3BGRL-2 revealed significant structural homology between the yeast protein fragment and the three human proteins (**Figure 1C**). All four structures share the characteristic Trx fold architecture, consisting of a central four-stranded β-sheet flanked by four α-helices, with isoform-specific structural differences confined to their respective C-termini.

### SH3BGRL proteins are monomeric in solution

Dimerization in solution was previously reported for human SH3BGRL-3 based on X-ray crystallographic studies (PDB-ID: 1T1V) and unpublished analytical size-exclusion chromatography experiments [9], while human SH3BGRL (PDB-ID: 1U6T) was reported to be monomeric in solution [10]. The C-terminal region of Aip5 is reported to form dimers in solution [27]. Assessment of the available crystal structures using the PDBePISA webserver does not support a dimeric state in solution [28]. As a precise knowledge about the oligomerization state in solution is of importance for our understanding of the SH3BGRL-actin interaction, we decided to systematically reanalyze the oligomerization state of the purified recombinant human SH3BGRL proteins. We produced the three human SH3BGRL isoforms recombinantly and purified the proteins to homogeneity (**Figure 2A**). Next, we performed sedimentation velocity experiments using analytical ultracentrifugation over a range of protein concentrations to determine the oligomerization state of the recombinant SH3BGRL isoforms (**Figure 2B**). SH3BGRL, SH3BGRL-2 and SH3BGRL-3 sedimented as single species with sedimentation coefficients s_20,w_ of 1.55 S, 1.49 S and 1.41 S, respectively. From the continuous c(s) distribution model in the program SEDFIT [29] molar masses of 13.0 kg mol^−1^, 12.4 kg mol^− 1^ and 10.1 kg mol^−1^, respectively, were obtained. These results agree very well with the masses calculated from the amino acid compositions and clearly support a monomeric state of all three isoforms under the chosen experimental conditions. From the molar masses and the sedimentation coefficients, frictional ratios f/f_0_ of 1.32, 1.30 and 1.27 can be calculated for SH3BGRL, SH3BGRL-2 and SH3BGRL-3, respectively, showing that all three isoforms deviate only moderately from the shape of a sphere.

**Figure 2:**
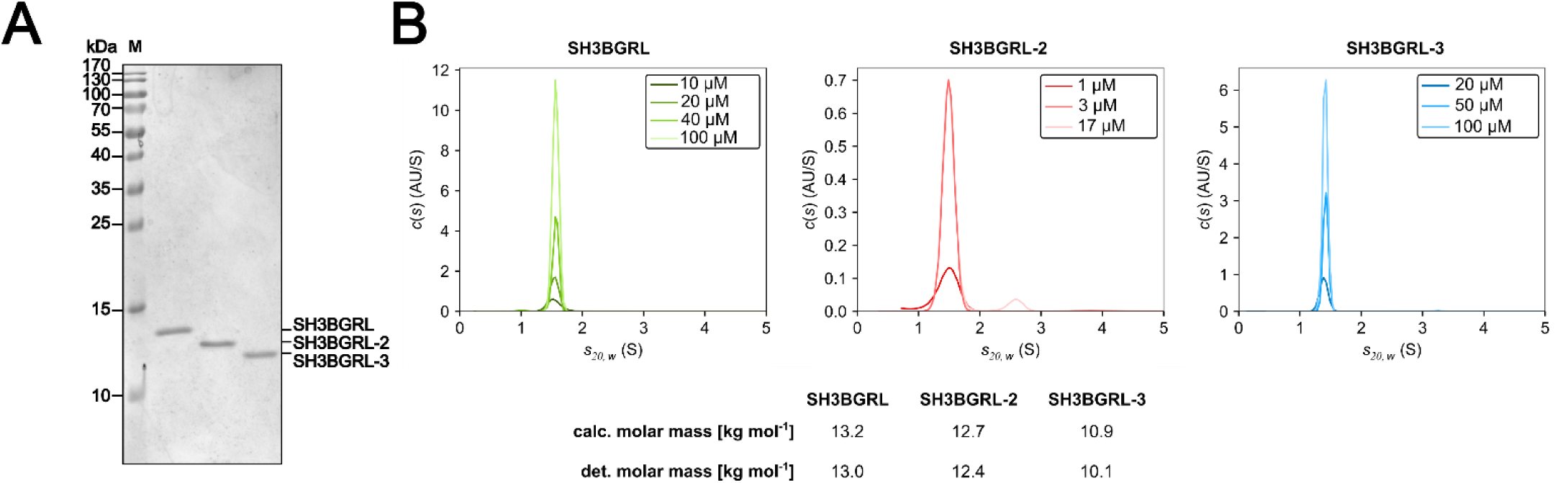
Analysis of the oligomerization state of human SH3BGRL proteins in solution. **(A)** SDS-PAGE of the purified recombinant human SH3BGRL isoforms used for biochemical studies. **(B)** Sedimentation velocity analyses of SH3BGRL, SH3BGRL-2 and SH3BGRL-3 at the indicated protein concentrations. The proteins sediment independently of the protein concentration as single species at s_20,w_=1.55 S (SH3BGRL), s_20,w_=1.49 S (SH3BGRL-2) and s_20,w_=1.41 S (SH3BGRL-3). The experimentally determined molar mass (det.) and the molar mass calculated from the amino acid composition (calc.) are given in the table. Note that the SH3BGRL-2 preparation contains a minor nucleic acid impurity that sediments at about 2.6 S. Because the sedimentation profiles at 1 µM and 3 µM were recorded at a wavelength of 230 nm and the profile at 17 µM was recorded at 280 nm, the minor nucleic acid impurity is only visible in the c(s) distribution at 17 µM.

### SH3BGRL proteins do not interact with G-actin

The C-terminal region of Aip5 has been reported to bind G-actin [27], and although the primary sequence of the identified responsible loop is not conserved in human SH3BGRL isoforms (**Figure 1A**), we decided to investigate whether monomeric human SH3BGRL isoforms can bind monomeric actin to form a heterodimeric complex.

We used AlphaFold3 and AlphaFold-Multimer to generate models of the potential SH3BGRL/- 2/-3 – G-actin complex [30]. We initially evaluated models generated by AlphaFold3 by plotting their respective predicted template modelling (pTM) and the interface predicted template modelling (ipTM) scores, which are metrics evaluating the accuracy of the predicted folding and the interaction interface, respectively. (**Figure 3A, B**). The pTM scores for all predictions ranged between 0.7 and 0.8, indicating a reliable prediction of the individual folds of the complex partners. In contrast, the ipTM scores revealed either failed or low-confidence predictions for the interactions within the complex. The most prominent complex predictions place the SH3BGRL proteins at the pointed end of the actin monomer (**Figure 3C**), closely resembling the arrangement of the penultimate actin protomer and SH3BGRL-2 in the available cryoEM-structure of the SH3BGRL-2 decorated actin filament [20] (**Suppl. Figure 1**). This indicates bias of AlphaFold3 towards this particular structural arrangement, likely due to inclusion of this structure in the training dataset or to the use of structural templates. To reduce this potential bias, we repeated the predictions using AlphaFold-Multimer, both with and without the use of templates (**Figure 3A, B**). These new predictions yielded a wider range of complex conformations. Under both conditions, AlphaFold-Multimer produced high-confidence predictions (ipTM > 0.8). However, most of these high-confidence predictions again placed SH3BGRL proteins at the pointed end of the actin monomer, similar to the conformations predicted by AlphaFold3. An exception was SH3BGRL-2, which was confidently positioned at the barbed end of the actin monomer in predictions using AlphaFold-Multimer with structural templates (**Figure 3C**).

**Figure 3:**
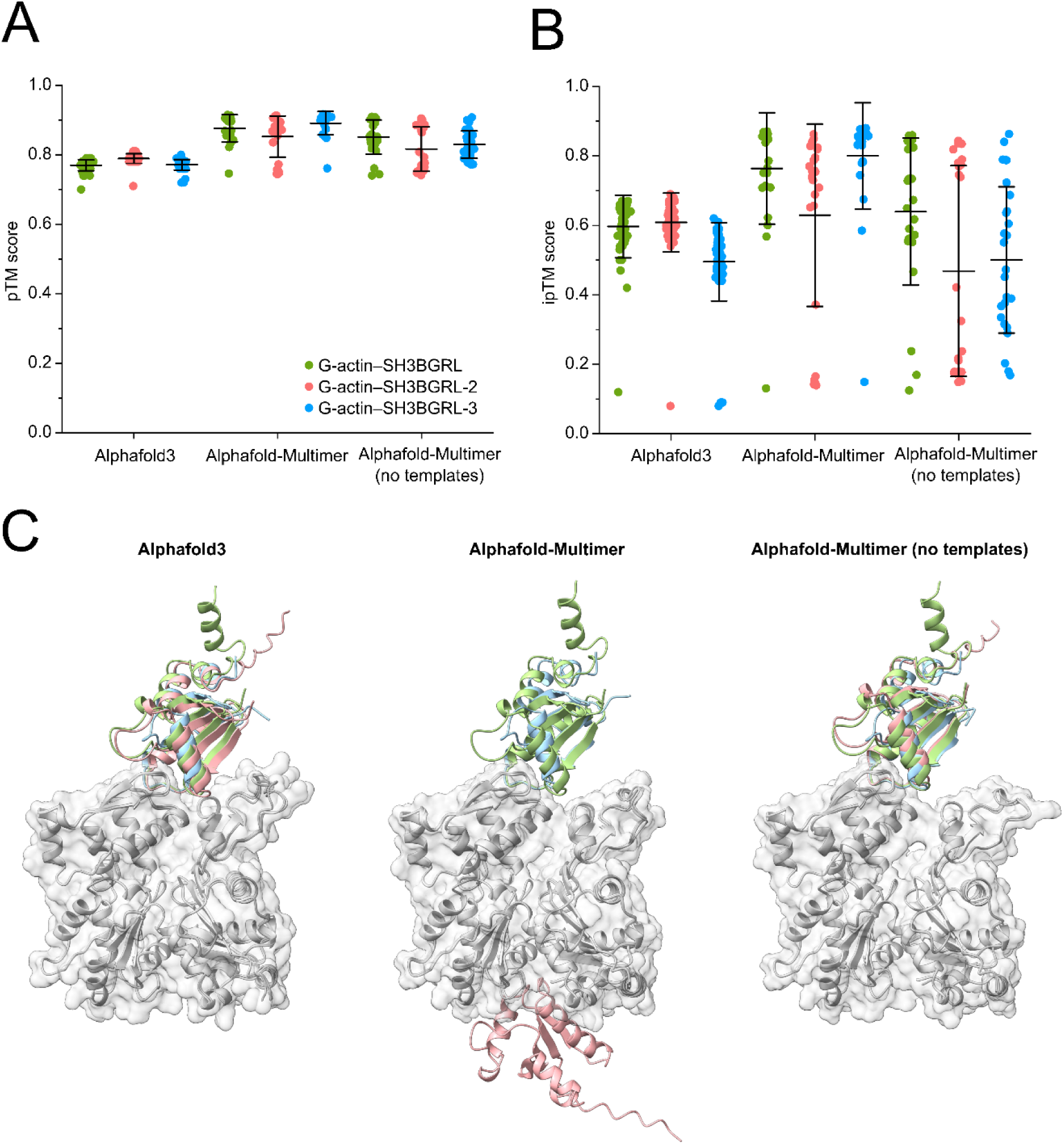
Investigation of potential SH3BGRL–G-actin complexes using AlphaFold. **(A)** Plotted predicted Template Modeling (pTM) scores for predictions of heterodimeric complexes of human SH3BGRL proteins with monomeric human β-actin derived using the indicated AlphaFold version. **(B)** Plotted interface predicted Template Modeling (ipTM) scores of the predictions shown in (A). **(C)** Overlay of the best ranked models of the SH3BGRL–G-actin complexes (SH3BGRL in green, SH3BGRL-2 in red, SH3BGRL-3 in blue, β-actin in grey) derived from the different used AlphaFold versions.

Motivated by the high discrepancy between the results obtained from different AlphaFold versions, we sought to investigate the potential SH3BGRL/-2/-3 – G-actin complex *in vitro*. G-actin binding proteins often modulate the exchange of the actin-bound nucleotide with the surrounding environment (e.g. profilins, thymosin β-4, cofilin) [31] (**Supplementary Figure 2**). We performed nucleotide dissociation experiments in the presence and absence of SH3BGRL-2 using the fluorescent ATP-analogue ɛ-ATP (**Figure 4A**) to infer possible binding of the isoform to G-actin. SH3BGRL-2 did not have any noticeable impact on the dissociation rate (*k*_-T_) of ɛ-ATP from the actin monomer. Next, we performed analytical size-exclusion chromatography experiments using G-actin and high molar excess of SH3BGRL-3 in the presence of latrunculin B (LatB), a marine toxin inhibiting actin polymerization, in G-buffer (**Figure 4B**). We did not observe a shift of the elution volume of G-actin in the presence of SH3BGRL-3, indicating the absence of a stable complex.

**Figure 4:**
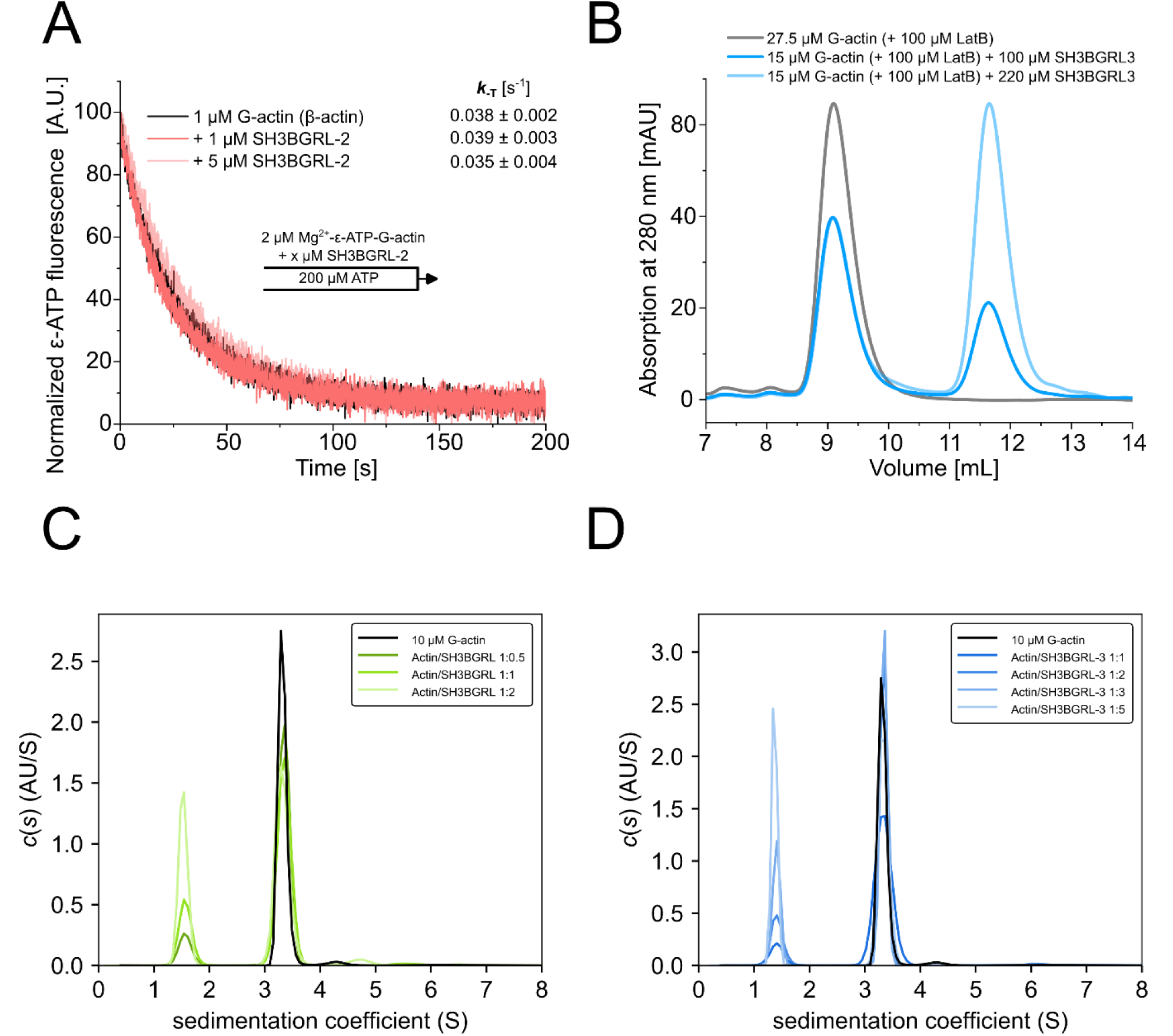
*In vitro* analysis of the potential SH3BGRL–G-actin complex. **(A)** The rate of nucleotide dissociation (*k*_-T_) from G-actin was determined for monomeric β-actin in the absence and presence of SH3BGRL-2 using fluorescently labeled ATP (ε-ATP) in G-buffer. Shown are representative experimental traces. The indicated values for *k*_-T_ are the mean ± SD of 11-12 individual mixing experiments and were determined by fitting mono-exponential functions to the experimental traces. **(B)** Analytical size exclusion chromatography experiments performed with LatB-stabilized G-actin in the absence and in the presence of a high molar excess of SH3BGRL-3 on a S75 10/300 size exclusion chromatography column (GE Healthcare, Chicago, USA) in G-buffer. No shift in the position of the G-actin peak was observed, indicating the absence of a stable SH3BGRL-3–G-actin complex. **(C, D)** Sedimentation velocity analysis performed with 10 µM LatB-stabilized G-actin in the absence and presence of increasing concentrations of SH3BGRL (C) or SH3BGRL-3 (D) under buffer conditions stimulating actin assembly (9 mM Tris pH 7.8, 50 mM KCl, 5 mM MgCl_2_, 0.18 mM CaCl_2_, 45 µM ATP, 0.18 mM TCEP). The sedimentation coefficient of the faster sedimenting LatB-stabilized G-actin (3.3 S) did not increase in the presence of the SH3BGRL isoforms, clearly indicating that both proteins do not interact with G-actin under these conditions.

To rule out the possibility of a weakly interacting complex undetectable by the methods used so far, we performed sedimentation velocity experiments with a constant concentration of 10 µM LatB-treated G-actin and increasing concentrations of SH3BGRL or SH3BGRL-3 under buffer conditions commonly used in biochemical studies of actin filament nucleation and elongation. The sedimentation coefficient of actin (3.3 S) did not change in the presence of SH3BGRL or SH3BGRL-3, indicating that SH3BGRL proteins and G-actin do not interact under these conditions (**Figure 4C, D**).

### The Trx fold of SH3BGRL isoforms forms a distinctive interface with the F-actin pointed end, with the C-terminus acting as a discriminator between SH3BGRL isoforms

In the available experimental structure of the spectrin–actin complex isolated from porcine erythrocytes, SH3BGRL-2 was identified as associated with the pointed end of the actin filament [20]. To comprehensively characterize the interaction between SH3BGRL isoforms and the F-actin pointed end, we performed all-atom molecular dynamics simulations. Initial models were generated by positioning AlphaFold3 models at the position of SH3BGRL-2 in the experimental structure (PDB: 8IAH) (**Figure 5A**). Throughout the simulation, the Trx fold of all SH3BGRL isoforms maintained its integrity and showed only minor deviations from the initial model position (**Figures 5B, 5C**). The simulation converged to a stable conformation after 350 ns, as indicated by the stable root-mean-square deviation (RMSD) from this point onwards (**Figure 5C**). The observed flexibility prior to convergence primarily originated from the C-terminal regions of SH3BGRL and SH3BGRL-2, which are absent in SH3BGRL-3 (**Figure 5D**).

**Figure 5:**
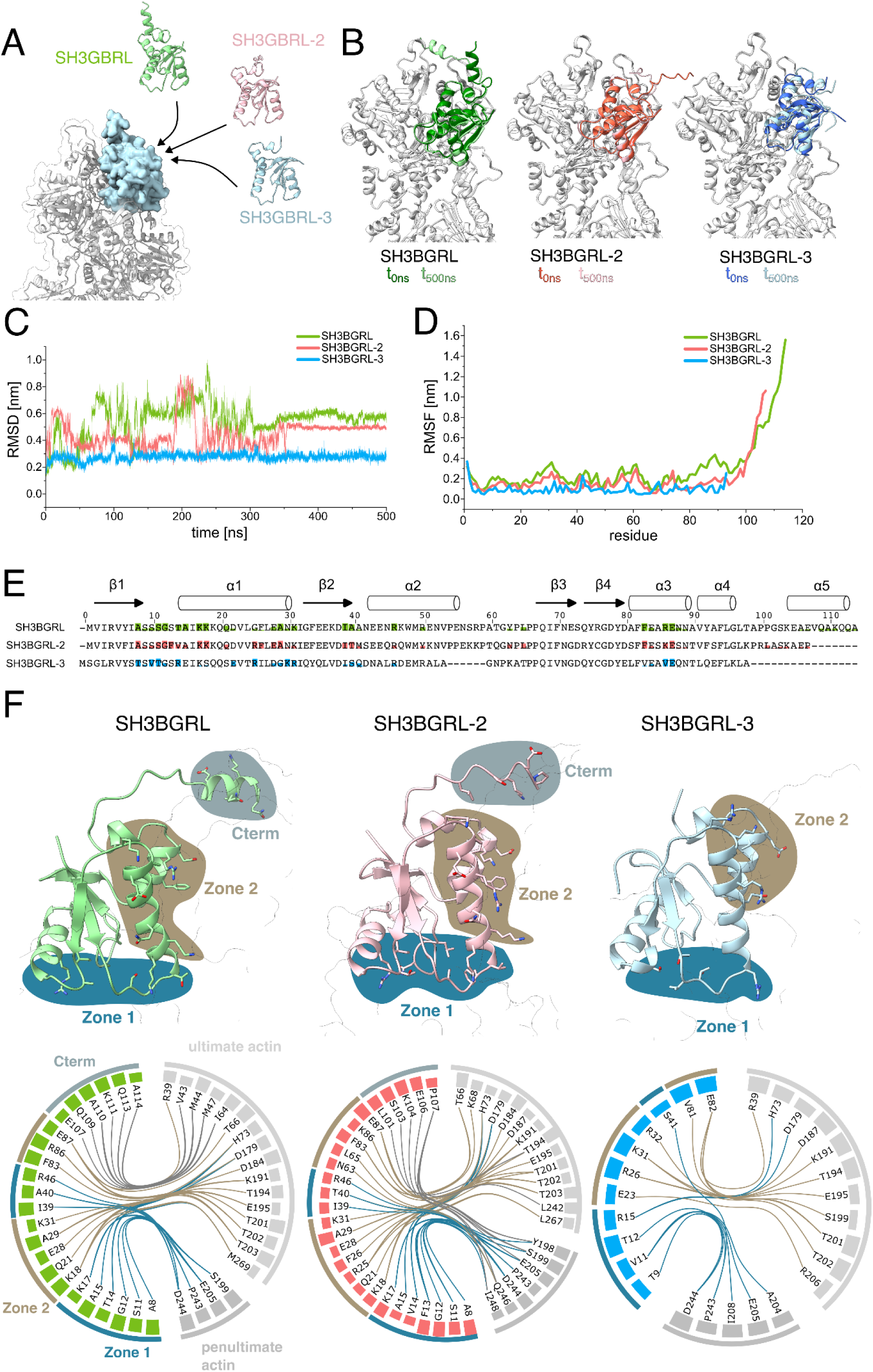
Analysis of the contact interface of SH3BGRL isoforms with the F-actin pointed end using all-atom molecular dynamics simulations. **(A)** The structures of human SH3BGRL isoforms (SH3BGRL – green, SH3BGRL-2 – red, SH3BGRL-3, blue) were predicted using AlphaFold and modeled into the position of SH3BGRL-2 at the F-actin pointed end in the experimental structure of the spectrin–actin complex (PDB-ID: 8IAH). **(B)** Overlay of the first (dark colors) and last frame (light colors) obtained from the molecular dynamics simulations at 0 ns and 500 ns, respectively. **(C)** RMSD values of the SH3BGRL isoforms along the simulation trajectory. **(D)** RMSF values of the SH3BGRL isoforms along the simulation trajectory. **(E)** Sequence alignment of SH3BGRL isoforms, where residues are filled with color according to the fraction of simulation time they spend in contact with actin. **(F)** Contact analysis of the last 100 ns of the simulation. Contacts are split into zones (Zone 1, Zone 2, Cterm) and are indicated by color. Upper panel: Models of SH3BGRL isoforms at the pointed end after 500 ns simulation, with contact residues shown as stick-style residues. Lower panel: Flareplots showing contacts that exist for at least 50% of the complete simulation time. Residues are shown as nodes and contacts as edges, with the colored bars at the individual residues indicating the relative contact frequency of that specific residue. Edges are colored using the color-scheme from the upper panel.

To identify key residues at the interaction interface between SH3BGRL isoforms and F-actin, we analyzed which residues made contact with actin throughout the simulation (**Supplementary Figure 3**). By combining this analysis with sequence alignment of the three isoforms, we correlated amino acid differences with variations in the binding interface. SH3BGRL, SH3BGRL-2 and SH3BGRL-3 showed similar interaction profiles, with the C-terminal stretch being the most significant difference (**Figure 5E, F**; **Supplementary Figure 3**).

For a more systematic characterization of the interaction interface, we divided the interaction profiles into three distinct zones (Zone 1, Zone 2, and Cterm) and visualized key interactions (present for >50% of the last 150 ns simulation time) between SH3BGRL isoforms and the F-actin pointed end (**Figure 5F**). The residues in Zone 1 mainly interact with the penultimate protomer. They involve the N-terminal part from A8 to K17 in SH3BGRL and SH3BGRL-2 and from T9 to R15 in SH3BGRL-3. Three additional residues (I39, A40/T40 and R46) between β-sheet 2 and α-helix 2 also contribute to Zone 1 and only bind to the penultimate protomer (**Figure 5F, Supplementary Figure 3**). In the SH3BGRL-3–F-actin complex, this contact is facilitated by S41. Only residue 15, located at the N-terminal tip of α-helix 1 represents a conserved contact to the ultimate protomer in Zone 1 in all three isoforms.

Zone 2 consists of residues located in α-helix 1 and α-helix 3, which contain predominantly charged residues (**Figure 5E, Supplementary Figure 3**). While SH3BGRL and SH3BGRL-2 show very similar interactions in this zone, SH3BGRL-3 shows fewer interactions. This is mainly due to a slight tilt in α-helix 1 that prevents the formation of contacts with actin at the N-terminal part of this helix (**Figure 5F**).

The C-terminal stretch (Cterm) of SH3BGRL and SH3BGRL-2 constitutes the third interaction motif. While the conserved Trx folds of SH3BGRL and SH3BGRL-2 interact very similar with the pointed end, this C-terminal stretch differs between the isoforms. For SH3BGRL, the C-terminal stretch interacts mainly with the last protomer, whereas for SH3BGRL-2 this region is 7 amino acids shorter, resulting in interactions mainly with the penultimate protomer (**Figure 5F**).

### All SH3BGRL isoforms interact with filamentous actin and inhibit actin depolymerization

Since all SH3BGRL isoforms appear to form stable complexes with the pointed end of an actin filament *in silico*, we wanted to assess their ability as actin end-binding proteins *in vitro*. We initially performed pyrene-actin based dilution-induced depolymerization studies with bare F-actin and F-actin pre-incubated with increasing concentrations of all three isoforms. We found that all three family members significantly reduced the observed depolymerization of pyrene-labeled actin filaments in a concentration-dependent manner (**Figure 6A**). Next, we performed depolymerization studies with F-actin capped at the barbed end by human capping protein (CP) to assess the potential pointed end capping observed in the published SH3BGRL-2–F-actin structure [20]. For this, we polymerized pyrene-labelled actin in the presence of CP and followed the depolymerization by diluting the sample below the critical concentration of the pointed end (∼ 0.6 µM). We initially tested our experimental setup by titrating human tropomodulin-3 (Tmod3), a well-characterized pointed end capping protein [32,33]. We observed a strong and concentration-dependent decrease in the observed rate of pointed end depolymerization, indicating association of Tmod3 with the pointed end (**Supplementary Figure 4A**), which allows us to estimate the dissociation constant (*K*_D_) of the Tmod3–pointed end complex to be 0.13 ± 0.02 µM (**Figure 6C**). This value is in good agreement with previous *K*_D_ values obtained using similar methods [32]. Next, we investigated the pointed end capping activities of the SH3BGRL proteins by titrating SH3BGRL and SH3BGRL-3. Both proteins showed a similar but significantly weaker effect on the pointed end depolymerization, when compared to Tmod3 (**Figure 6B**), which result in estimated *K*_D_ values of 7.58 ± 0.94 µM (SH3BGRL) and 7.68 ± 0.97 µM (SH3BGRL-3), respectively (**Figure 6C**). To assess whether SH3BGRL, the longest isoform, and Tmod3 can occupy the pointed end simultaneously, as shown for the SH3BGRL-2 [20], we performed experiments with a constant concentration of Tmod3 (0.375 µM) and SH3BGRL. We found that both proteins together inhibited pointed end depolymerization stronger than individually, indicating that they do not displace each other at the pointed end (**Supplementary Figure 4B**). An overlay of the last frame of the SH3BGRL– F-actin MD simulation and the published spectrin–actin structure, which also contains tropomodulin, supports this notion. Although the C-terminal part of SH3BGRL is in close proximity to tropomodulin, no major steric hindrances are expected, also due to the flexibility of this region observed in the MD simulations (**Supplementary Figure 4C**).

**Figure 6:**
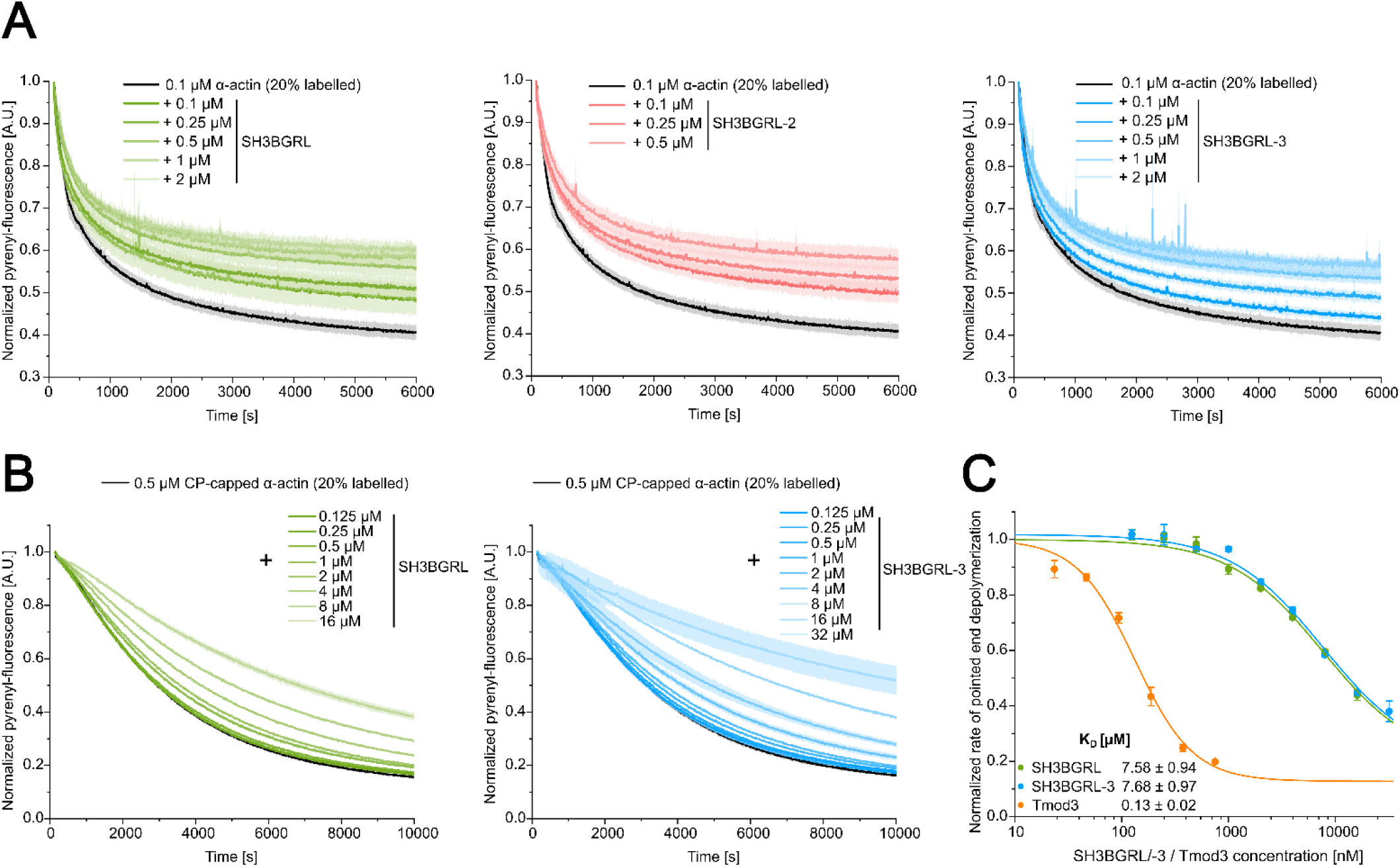
Analysis of the pointed end capping activity of SH3BGRL isoforms using pyrene-actin based depolymerization experiments. **(A)** Pyrene-actin based dilution-induced depolymerization experiments the absence and presence of increasing concentrations of SH3BGRL isoforms were used to examine the effect of the isoforms on filament depolymerization. Data are shown as the mean ± SD. For each condition at least N=3 individual experiments were performed. **(B)** Pyrene-actin based dilution-induced depolymerization experiments using filaments capped at the barbed end by human capping protein (CP) were used to specifically monitor the effect of SH3BGRL isoforms on pointed end depolymerization. Data are shown as the mean ± SD. N=3 for each condition. **(C)** The binding affinities (K_D_) of SH3BGRL/-3 and Tmod3 to the pointed end were estimated by plotting the normalized pointed end depolymerization rates against the used protein concentration and fitting a hyperbolic function to the complete dataset. The traces of the depolymerization experiments performed in the presence of Tmod3 are shown in Supplementary Figure 4.

### SH3BGRL isoforms stimulate actin polymerization from G-actin but not from profilin– G-actin

Our analytical ultracentrifugation studies indicate that SH3BGRL proteins do not directly interact with G-actin in solution under conditions that stimulate actin assembly. Furthermore, all isoforms interact with F-actin as shown by our *in silico* analysis and *in vitro* depolymerization studies. Therefore, we hypothesized that SH3BGRL proteins might influence actin polymerization initiated from G-actin by interacting with newly formed oligomers and filaments. To challenge our hypothesis, we performed pyrene-actin based bulk-polymerization assays with pyrene-labeled α-actin and increasing concentrations of human SH3BGRL and SH3BGRL-3 isoforms (**Figure 7A**). These experiments demonstrated a concentration-dependent stimulatory effect of both isoforms on actin polymerization. Specifically, they shortened the initial lag time of the reaction and enhanced the subsequent linear increase in pyrenyl-fluorescence, which is typically correlated with the net polymerization rate of actin filaments, up to 3-fold **(Figure 7B, C**). We repeated the experiments using a specific concentration of SH3BGRL-2, which demonstrated a similar but less pronounced effect on actin assembly. *In vivo*, most of G-actin is sequestered by thymosin-β4 or bound to members of the profilin protein family. The profilin–G-actin complex constitutes the predominant polymerization-competent actin species in the cytosol, which can be used by actin polymerases like formins to nucleate and elongate filaments [34]. We therefore analysed if the SH3BGRL isoforms can stimulate actin polymerization started from profilin–G-actin by repeating our assays in the presence of an excess of profilin-1. A nanomolar concentration of diaphanous-related formin-1 (mDia1) is able to overcome profilin-induced inhibition of spontaneous actin polymerization (**Figure 7D**). Conversely, polymerization traces derived from experiments performed with profilin–G-actin in combination with SH3BGRL isoforms are indistinguishable from those obtained with profilin–G-actin alone, demonstrating that SH3BGRL proteins are unable to counteract profilin-induced inhibition of actin polymerization (**Figure 7D**).

**Figure 7:**
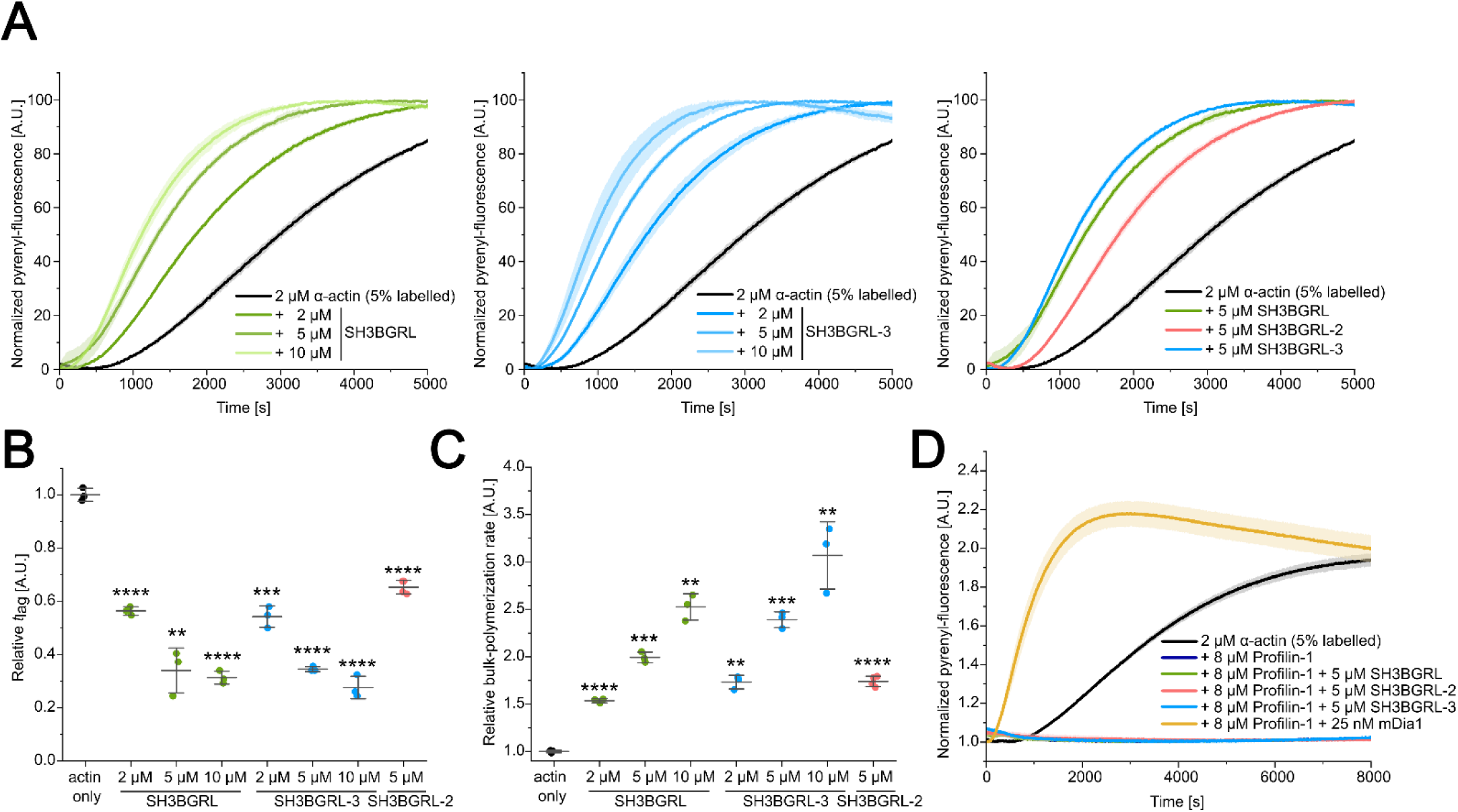
Analysis of the effect of SH3BGRL isoforms on actin polymerization started from G-actin and profilin–G-actin using pyrene-actin based bulk-experiments. **(A)** Pyrene-based polymerization experiments with 2 µM α-actin (5% pyrene-labeled) in the absence and presence of different concentrations of SH3BGRL isoforms. The solid lines/shades represent the mean ± SD of three individual experiments. **(B)** Relative values of *t*_lag_ (to actin only experiments) determined from experiments shown in (A). *t*_lag_ was determined as the time-point at which the reaction reaches 5% of the final fluorescence signal. Data are shown as the mean ± SD of all performed experiments. N=3 for each condition. Significance is given compared to actin only experiments (p > 0.05 ≙ ns, p ≤ 0.05 ≙ *, p ≤ 0.01 ≙ **, p ≤ 0.001 ≙ ***, p ≤ 0.0001 ≙ ****) **(C)** Relative bulk-polymerization rates (to actin only experiments) determined from the experiments shown in (A). Data are shown as the mean ± SD of all performed experiments. N=3 for each condition. Significance is given compared to actin only experiments (p > 0.05 ≙ ns, p ≤ 0.05 ≙ *, p ≤ 0.01 ≙ **, p ≤ 0.001 ≙ ***, p ≤ 0.0001 ≙ ****). **(D)** Pyrene-based polymerization experiments with 2 µM α-actin (5% pyrene-labeled) in the presence of 8 µM profilin-1 and SH3BGRL/-2/-3 or mDia1. The solid lines/shades represent the mean ± SD of three individual experiments.

### SH3BGRL proteins stimulate filament nucleation but do not affect filament elongation

Actin polymerization can be described as a two-step process in which a thermodynamically and kinetically unfavorable nucleation step, the formation of an actin-trimer, is followed by a more favorable asymmetric elongation of the trimer into an actin filament [35]. Pyrene-actin based bulk-experiments allow no clear discrimination of these two distinct processes. Therefore, to get a better understanding of the underlying mechanism of SH3BGRL-stimulated actin assembly, we performed single-filament assays with ATTO-488 labeled α-actin using total internal reflection fluorescence microscopy (TIRFM).

We analyzed the effect of SH3BGRL-3 on actin polymerization in single-filament assays by titrating a constant concentration of monomeric ATTO-488 labeled α-actin (1 µM) with increasing concentrations of SH3BGRL-3 in G-buffer. Actin polymerization was induced by salt-shift, the reaction mixture flushed into a flow-cell and imaging was initiated. SH3BGRL-3 significantly increased the filament density over the entire duration of the experiments in a concentration-dependent manner compared to experiments performed without SH3BGRL-3 (**Figure 8C**). We determined the apparent rate of filament nucleation (*k*_nuc_) by linear regression analysis of the linear regions of the filament density time course (**Figure 8A**, **Table 1**). At the highest concentration used (5 µM), SH3BGRL-3 increased *k*_nuc_ 6.7-fold compared to experiments performed with actin alone. We repeated the experiments with selected concentrations of SH3BGRL and SH3BGRL-2 (**Figure 8A, C**). While SH3BGRL showed comparable effects on *k*_nuc_, the stimulating effect of SH3BGRL-2 on actin nucleation was significantly lower, which is in line with the results obtained from the pyrene-actin bulk-experiments. Next, we tracked the growth of individual actin filaments under the different experimental conditions to investigate the effect of the SH3BGRL proteins on filament elongation. We observed no significant effect of all three isoforms on the apparent rate of filament elongation (**Figure 8B**). Most of our biochemical studies to this point were performed using α-actin from chicken pectoralis major muscle, as this isoform is easily obtained directly from tissue with high purity and yield and is routinely used in studies of actin dynamics. To assess whether SH3BGRL isoforms can also modulate actin dynamics of other, physiologically more relevant isoforms, we repeated our TIRFM-based experiments using purified recombinant human cytoskeletal β-actin and a selected concentration of SH3BGRL-3 (1 µM). We observed a similar but stronger stimulation of actin nucleation compared to experiments performed with α-actin and the same concentration of SH3BGRL-3 and, in agreement with our previous experiments, no effect on filament elongation (**Supplementary Figure 5**). In conclusion, our data indicates that SH3BGRL proteins stimulate actin assembly solely by increasing the rate of filament nucleation, most likely by stabilizing energetically unstable actin dimers and trimers.

**Figure 8:**
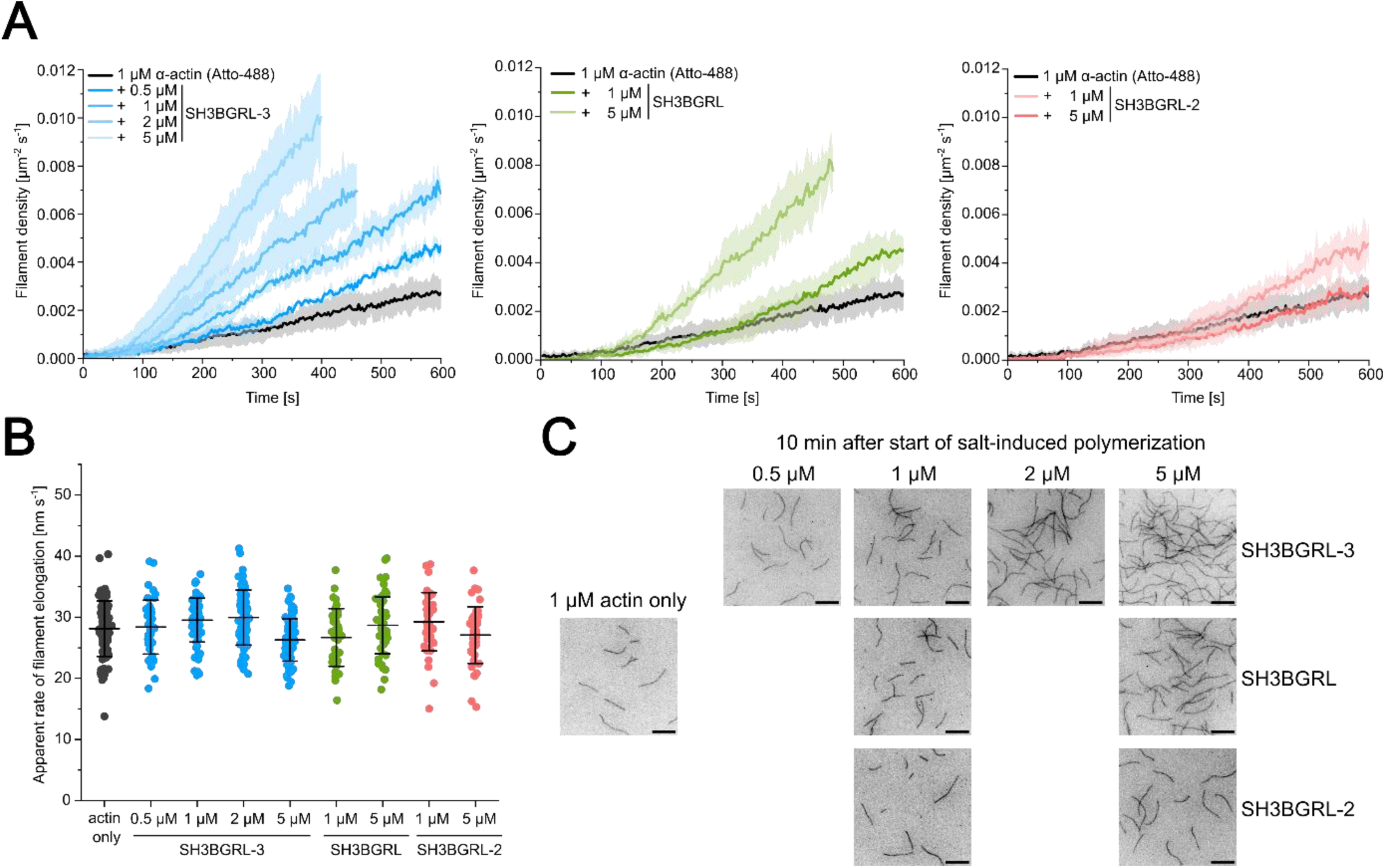
Analysis of the effect of SH3BGRL isoforms on actin nucleation and elongation using *in vitro* TIRF microscopy. **(A)** Polymerization of 1 µM fluorescently labeled α-actin (15% Atto-488 labeled) was induced by salt-shift in the absence or presence of different concentrations of SH3BGRL isoforms and the progression of the reaction was tracked by TIRF microscopy. The increase in filament density over time was tracked using the *Analyze particles* plugin in ImageJ. The solid lines and shades represent the mean ± SD of at least three individual experiments. Nucleation rates were determined by fitting linear functions to the regions of linear increase. **(B)** The elongation rates of individual filaments were determined by manual tracking of the elongating filaments. Every data point represents an individual filament. Data is shown as the mean ± SD. No significant changes were observed among the various conditions. **(C)** Representative micrographs of TIRFM-based polymerization experiments under the indicated conditions 10 min after induction of polymerization. Scale bar corresponds to 10 µm.

**Table 1:** Actin filament elongation and nucleation rates in the absence and presence of SH3BGRL/-2/-3 determined from TIRFM-based experiments.

| Parameter | SH3BGRL | SH3BGRL-2 | SH3BGRL-3 |
| --- | --- | --- | --- |
| Apparent rate of filament elongation [nm s<sup>-1</sup>] | 28.1 ± 4.5 (actin only)<br>-<br>26.7 ± 4.7 (1 μM)<br>-<br>28.7 ± 4.6 (5 μM) | 28.1 ± 4.5 (actin only)<br>-<br>29.3 ± 4.8 (1 μM)<br>-<br>27.1 ± 4.6 (5 μM) | 28.1 ± 4.5 (actin only)<br>28.4 ± 4.4 (0.5 μM)<br>29.5 ± 3.6 (1 μM)<br>29.9 ± 4.5 (2 μM)<br>26.3 ± 3.5 (5 μM) |
| Apparent rate of filament nucleation $k_{\text{nuc}}$ [μm<sup>-2</sup> s<sup>-1</sup>] | 5.1×10<sup>-6</sup> ± 1.1×10<sup>-6</sup> (actin only)<br>-<br>12.3×10<sup>-6</sup> ± 1.2×10<sup>-6</sup> (1 μM)<br>-<br>22.2×10<sup>-6</sup> ± 3.0×10<sup>-6</sup> (5 μM) | 5.1×10<sup>-6</sup> ± 1.1×10<sup>-6</sup> (actin only)<br>-<br>7.0×10<sup>-6</sup> ± 1.2×10<sup>-6</sup> (1 μM)<br>-<br>12.0×10<sup>-6</sup> ± 1.4×10<sup>-6</sup> (5 μM) | 5.1×10<sup>-6</sup> ± 1.1×10<sup>-6</sup> (actin only)<br>10.5×10<sup>-6</sup> ± 0.8×10<sup>-6</sup> (0.5 μM)<br>13.8×10<sup>-6</sup> ± 2.5×10<sup>-6</sup> (1 μM)<br>18.4×10<sup>-6</sup> ± 3.0×10<sup>-6</sup> (2 μM)<br>34.2×10<sup>-6</sup> ± 6.0×10<sup>-6</sup> (5 μM) |

## Discussion

The thioredoxin (Trx) fold is among the most prevalent structural motifs in proteins, primarily known for its enzymatic roles. Numerous protein families featuring the Trx fold display variations of the classic catalytic motifs, CXXC and CXXS/T, which are key determinants of their enzymatic activity [2]. However, the SH3BGRL protein family, predominantly characterized by the Trx fold, is devoid of a catalytic motif, prompting inquiry into its functional significance. In this study, we explored the interactions between the universally produced SH3BGRL isoforms (SH3BGRL, SH3BGRL-2, SH3BGRL-3) and actin to unveil a potential role for this protein family in modulating actin dynamics.

Previous biochemical and structural characterizations of SH3BGRL proteins presented ambiguities regarding their oligomerization state. Through analytical ultracentrifugation, we have now demonstrated that all members of the SH3BGRL family exist in the micromolar concentration range as monomers in solution. This finding clarifies previous crystallography-based assumptions that suggested oligomerization for SH3BGRL-3 [9]. Our findings are consistent with the published monomeric structure of SH3BGRL-2 at the pointed end of actin filaments [20], reinforcing the hypothesis that all SH3BGRL isoforms interact with actin through a similar mechanism.

Historically, SH3BGRL isoforms have been considered molecular adaptor proteins, with their expression levels linked to cancer progression [12–15]. The naming of these proteins stems from the presence of a singular SH3-binding motif in the primary sequence of SH3BGRL and SH3BGRL-2, absent in SH3BGRL-3. However, both our structural analysis and previous studies reveal that this motif is sequestered within the tertiary structures of SH3BGRL and SH3BGRL-2, rendering direct interactions with SH3 domains improbable [9]. Furthermore, no experimental evidence supports direct binding of SH3BGRL proteins to SH3 domains. In light of these observations, the name of the SH3BGRL protein family may be misleading, and caution should be exercised when considering its functional implications.

Through a combination of direct and indirect experimental approaches, we demonstrated that SH3BGRL proteins lack G-actin binding properties. This finding contrasts with the results obtained for the C-terminal region of the actin-binding protein YFR016c/Aip5 from *Saccharomyces cerevisiae*, which, like SH3BGRL proteins, is predominantly composed of a Trx fold [27,36,37]. Notably, YFR016c/Aip5 relies on dimerization to enhance its actin nucleation capability, a characteristic absent in SH3BGRL proteins. Despite the lack of G-actin binding by SH3BGRL proteins, we observed that all SH3BGRL isoforms interact with the pointed end of actin filaments, resulting in reduced filament depolymerization in bulk-disassembly experiments. Additionally, we observe nucleating properties in both bulk-polymerization assays and single-filament experiments, but no detectable effect on barbed end elongation. Based on these observations, we propose that SH3BGRL proteins facilitate actin nucleation by stabilizing thermodynamically and kinetically unfavorable actin dimer and trimer states [35] in a structural arrangement reminiscent of the arrangement of SH3BGRL proteins at the pointed end of the filament, thereby allowing normal barbed end elongation from SH3BGRL-nucleated filaments.

Using molecular dynamics simulations followed by comprehensive contact analysis, we characterized the interaction between the Trx fold and the pointed end of the actin filament across all three isoforms. While the interaction site is not conserved at the amino acid sequence level, it is preserved structurally. This suggests that the Trx fold is the critical determinant for the interaction with the F-actin pointed end. The variations at the C-terminus represent the most significant differences among the isoforms, yet their influence on pointed end binding remains unclear. When comparing the nucleating effect on actin assembly of the three isoforms, SH3BGRL-2 shows the lowest potency. This seems to contrast with the interaction interface analysis, which shows a very similar interaction interface of SH3BGRL and SH3BGRL-2 with the pointed end. A possible explanation for this discrepancy could be the role of the C-terminal stretch, which is different in the two isoforms. A greater flexibility of this stretch in the SH3BGRL-2 isoform may hinder initial actin binding and thereby reduce the potency of actin nucleation.

Unlike well-characterized actin nucleators, SH3BGRL proteins lack the Wiskott-Aldrich homology 2 (WH2) domain, a ∼17-amino acid actin-binding motif commonly found in potent actin nucleators [38]. WH2 domains, often present in tandem repeats, are crucial for actin monomer recruitment and nucleation. Examples of WH2-containing nucleators include formins (e.g., FMNL3, mDia1), nucleation-promoting factors of the Arp2/3 complex (e.g., N-WASP, WAVE2) [39], SPIRE [40], Cordon Bleu [41], and the efficient actin nucleator and pointed end capping protein in muscle cells, Leiomodin [42]. Despite the prevalence of WH2 domains among efficient actin nucleators, selected proteins, such as tropomodulin isoforms, can also nucleate actin in their absence, suggesting alternative mechanisms of nucleation [43]. The same study showed that the nucleating properties of selected tropomodulin isoforms can be attributed, at least in part, to their G-actin binding. While tropomodulins and SH3BGRL proteins do not share G-actin binding properties, they have in common that they are relatively weak actin nucleators compared to potent WH2-containing nucleators, which exhibit highly efficient nucleation in the low nanomolar concentration range.

In the published structure of the spectrin–actin complex, SH3BGRL-2, along with Tmod1, is positioned at the pointed end of the actin filament. [20]. Our *in vitro* and *in silico* experiments with SH3BGRL and Tmod3, coupled with analysis of existing data on the structural configuration at the pointed end, suggest that the longest isoform can similarly occupy the point end in conjunction with tropomodulin. Therefore, we suggest that this is a common feature among the SH3BGRL protein family. How the Tmod and SH3BGRL protein families work together to regulate actin dynamics in the cellular context is a question that needs to be addressed in future studies.

SH3BGRL proteins are produced ubiquitously, while expression of the *SH3BGR* gene, which encodes the SH3BGR protein, the eponymous member of the protein family, is enriched in skeletal and cardiac muscle cells. [4,5]. Experimental evidence on the functional role of SH3BGR in muscle cells is rare. A recent study reported upregulation of SH3BGR in the failing heart of patients suffering from hypertrophic cardiomyopathy, as well as in murine models of the condition [11]. Furthermore, artificial overexpression in neonatal rat ventricular cardiomyocytes results in the upregulation of prominent HCM-markers [11]. An earlier study investigated the localization pattern of the SH3BGR protein in the *Xenopus leavis* heart muscle and found specific localization of the protein to the Z-line of the developed sarcomere [5]. In addition, knock-down of SH3BGR resulted in severely disrupted sarcomere formation. Localisation to the Z-line is inconsistent with the pointed end binding demonstrated in our *in vitro* experiments involving the ubiquitously produced SH3BGRL proteins. Similar to SH3BGRL isoforms, SH3BGR also contains a Trx fold domain and one SH3-binding/Homer EVH1 binding motif. The C-terminal region strongly deviates from SH3BGRL isoforms as it contains a long C-terminal extension rich in glutamic acid residues [6–8]. Future *in vitro* and *in vivo* studies will need to investigate the potential role of the SH3BGR protein in sarcomere formation and stabilisation in the context of our present work and delineate a possible role of the C-terminal extension, which may explain the observed localisation pattern.

In conclusion, our study provides key mechanistic insights into the involvement of the SH3BGRL protein family in actin dynamics and sheds light on an expanded function of the thioredoxin fold beyond its conventional enzymatic function, demonstrating its potential as a critical regulatory component in cytoskeletal organization.

## Material and Methods

### Plasmids

The coding sequences of human SH3BGRL (Uniprot-ID: O75368), human SH3BGRL-2 (Uniprot-ID: Q9UJC5) and human SH3BGRL-3 (Uniprot-ID: Q9H299) were obtained from the Uniprot database, and the corresponding cDNA sequences were synthesized using the GeneArt Gene Synthesis service (Thermo Fisher Scientific, Waltham, MA, USA). The coding sequences were cloned into the pGEX-6P-2 vector for expression as GST-fusion proteins and verified via overnight sequencing. The coding sequence of human cytoskeletal β-actin (Uniprot-ID: P60709) was fused via a C-terminal linker (ASR(GGS)_3_A) to a His_8_-tagged thymosin-β4 moiety (Uniprot-ID: P62328) and cloned into the multiple cloning site of the pFastBac-Dual vector under the control of the polyhedrin promotor.

### Protein production and purification

Human GST-SH3BGRL/-2/-3 fusion proteins were produced in Rosetta2 cells. For a typical preparation, cells from 1 liter of expression culture were resuspended in 100 mL lysis buffer (25 mM HEPES pH 7.4, 150 mM NaCl, 15 mM CaCl_2_, 1 mM DTT, 1 mM PMSF, 100 µg/mL TAME, 80 µg/mL TPCK, 2 µg/mL pepstatin, 5 µg/mL leupeptin) supplemented with 250 µg/mL lysozyme (from hen egg white; Merck KGaA, Darmstadt, Germany) and incubated on ice for 30 minutes. The cells were lysed by sonification and treated with DNase-I (Roche, Basel, Switzerland) for 30 minutes on ice to remove bacterial DNA. The lysate was cleared by centrifugation at 35,000 × *g* for 30 minutes and the cleared supernatant was subsequently loaded onto a self-packed GSH-Sepharose column and washed with 5 column volumes of wash buffer (25 mM HEPES pH 7.4, 150 mM NaCl, 1 mM DTT, 1 mM EDTA). The fusion protein was digested on-column with PreScission protease overnight at 4°C. On the next day the SH3BGRL/-2/-3 was eluted from the column with wash buffer. The protein was concentrated and loaded onto a S75 16/600 size exclusion chromatography column (GE Healthcare, Chicago, IL, USA) equilibrated with SEC-buffer (25 mM HEPES pH 7.4, 50 mM NaCl, 0.5 mM TCEP). Fractions containing pure protein were pooled, concentrated, frozen in liquid nitrogen and stored at −80 °C until used further. Purification of human cytoskeletal β-actin and labelling at the accessible C-terminal cysteine with ATTO-655 (ATTO-TEC, Siegen, Germany) was performed exactly as previously described [44]. α-skeletal actin (UniProt ID: P68139) was prepared from chicken pectoralis major muscle and labeled at the accessible C-terminal cysteine with *N*-(1-pyrene)iodoacetamide (Thermo Fisher Scientific, Waltham, MA, USA) or ATTO-488 (ATTO-TEC, Siegen, Germany) following previously described protocols [45–47]. To further purify actin for use in polymerization studies and for analytical ultracentrifugation, the crude G-actin solution was centrifuged (186,000 × *g*, 2 hours, 4°C) and loaded onto a S75 16/600 size exclusion chromatography column (GE Healthcare, Chicago, IL, USA) equilibrated with G-buffer (10 mM TRIS pH 8.0, 0.2 mM CaCl_2_, 0.2 mM ATP, 0.5 mM DTT). Only fractions from the second half of the chromatography peak were used for polymerization studies and analytical ultracentrifugation. Production and purification of human thymosin-β4-His_8_ [44], human cofilin-1 [48], human profilin-1 [48], mouse SNAP-mDia1(FH1FH2)-His_6_ [49] and human tropomodulin-3 [50] were performed as previously described.

### Nucleotide exchange assay

The rate of ATP dissociation (*k*_-T_) for monomeric cytoskeletal β-actin in the absence and presence of SH3BGRL-2, profilin-1, thymosin-β4-His_8_ and cofilin-1 was determined using ε-ATP (Jena Bioscience, Jena, Germany) at a HiTech Scientific SF61 stopped-flow system (TgK Scientific Limited, Bradford on Avon, UK), as previously described [48]. For experiments in the presence of actin-binding proteins (ABPs), actin and the respective ABP were incubated on ice for 15 min prior to experiment.

### Analytical size-exclusion chromatography

Freshly purified monomeric skeletal α-actin was treated with a 1.5 molar excess of latrunculin B (LatB) for use in analytical size-exclusion chromatography. LatB-actin and SH3BGRL-3 were diluted into G-buffer to the indicated concentrations and a final volume of 100 µL and incubated on ice for 15 min. The mixture was injected onto a S75 10/300 size-exclusion chromatography column (GE Healthcare, Chicago, IL, USA) equilibrated with G-buffer (10 mM TRIS pH 8.0, 0.2 mM CaCl_2_, 0.5 mM DTT).

### Pyrene-actin based bulk-polymerization assays

Pyrene-actin based bulk-polymerization experiments in the absence and presence of SH3BGRL proteins were performed at a CLARIOstar Plus microplate reader (BMG LABTECH, Offenburg, Germany) at 25 °C. Pyrene-labeled Mg^2+^-ATP-G-actin (5% labeled) was premixed with SH3BGRL proteins. 20 µL of the individual reaction mixtures were placed in a black flat-bottom 96-well plate (BrandTech Scientific, USA). 80 μL of 1.25× polymerization buffer (10 mM HEPES pH 7.4, 50 mM KCl, 5 mM MgCl_2_, 0.5 mM EGTA, 0.1 mM DTT, and 0.1 mM ATP) was added to the wells using the built-in pipetting function of the plate reader, and the polymerization of actin was followed as an increase in pyrenyl-fluorescence. For experiments in the presence of profilin-1 and mDia1, all components were premixed on ice and then placed in the 96-well plate. The bulk-polymerization rates were determined by linear regression analysis of the linear region around the time-point of half-maximal fluorescence [51]. The lag-time (*t*_lag_) of the reaction was defined as the time-point at which the reaction reaches 5 % of the final fluorescence signal. For dilution-induced depolymerization experiments, 5 µM of pre-polymerized pyrene-labeled F-actin (20% labeled) was incubated with SH3BGRL proteins on ice for 15 min. The reaction mixtures were rapidly diluted to 0.1 µM by addition of polymerization buffer using the same approach as described for the polymerization assays. The dilution-induced depolymerization of actin filaments was followed as a decrease in pyrenyl-fluorescence. For depolymerization experiments specifically probing the pointed end, the protocol was slightly modified. Pyrene-labeled actin (10 µM, 20% labeled) was polymerized overnight in the presence of 0.1 µM human capping protein (CP) to generate CP-capped actin filaments. 10 µM of pre-polymerized pyrene-labeled CP-capped F-actin (20% labeled) was incubated with SH3BGRL proteins on ice for 15 min. The reaction mixtures were rapidly diluted to 0.5 µM and the depolymerization of the filaments from the pointed ends was tracked via the decrease in pyrenyl-fluorescence. Mono-exponential functions were used to determine the apparent rate of pointed end depolymerization under the various conditions. The normalized depolymerization rates were plotted against the protein concentration used to estimate the respective *K*_D_ values via a hyperbolic function:

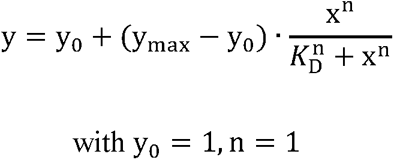

### TIRF microscopy-based assays

The effect of SH3BGRL proteins on filament nucleation and elongation was assessed using TIRF microscopy-based assays with freshly clarified ATTO-488 labeled α-actin and ATTO-655 labeled cytoskeletal β-actin. Glass coverslips used for flow cell assembly were cleaned and chemically treated according to previously described protocols [48]. Actin polymerization was initiated by diluting the G-actin solution (containing 15 % labeled actin (α-actin) or 10 % labeled actin (β-actin)) to a final concentration of 1 µM in 2×TIRF-buffer (20 mM HEPES, 50 mM KCl, 1 mM MgCl_2_, 1 mM EGTA, 0.2 mM ATP, 15 mM glucose, 20 mM β-ME, 0.25 % methylcellulose, 0.1 mg/mL glucose oxidase and 0.02 mg/mL catalase). After mixing, the solutions were immediately flushed into flow cells and image acquisition was started. For experiments involving SH3BGRL isoforms, the isoforms were pre-diluted in KMEI buffer (10 mM imidazole pH 7.4, 50 mM KCl, 1 mM MgCl_2_, 1 mM EGTA) and further diluted to the final concentration in TIRF-buffer before the addition of actin.

Image series were acquired at an Olympus IX83 inverted fluorescence microscopes (Olympus, Hamburg, Germany) equipped with a 60×/1.49 NA PlanApo TIRF oil immersion objective and an Orca Flash 4.0 CMOS camera (Hamamatsu Photonics Deutschland GmbH, Herrsching, Germany). Nucleation of actin filaments was analyzed by automated image analysis using the *Analyze particles* plugin in ImageJ [52]. The elongation rates of individual actin filaments were measured by manually tracking their growth.

### AlphaFold

AlphaFold3 [30] and AlphaFold-Multimer (v3) were used to predict potential heterodimeric complexes of human SH3BGRL proteins and human cytoskeletal β-actin. The respective primary sequences were retrieved from the Uniprot database (see **Plasmids**). Predictions using AlphaFold3 were performed using the AlphaFold server (https://golgi.sandbox.google.com). We ran 10 jobs on random-seed for each complex combination resulting in 50 models in total for each combination. The metrics for further evaluation of the complexes were extracted from the corresponding JSON files. Predictions using AlphaFold-Multimer (v3) with and without templates were performed on the Halime HPC cluster at the Institute for Biophysical Chemistry (Hannover Medical School, Germany). We generated a total of 25 complex predictions for each complex combination. Metrics for further analysis were extracted from the AlphaFold pickle files of each respective prediction via custom python scripts.

Visualization of the predicted complex structures was performed using ChimeraX [53].

### Molecular dynamics (MD) simulations

The individual complexes of the F-actin pointed end with the different SH3BGRL isoforms were modelled by positioning the AlphaFold3 generated [30] models of the SH3BGRL isoforms in the binding orientation observed for SH3BGRL-2 in the experimental structure of the spectrin–actin complex (PDB: 8IAH). Since SH3BGRL isoforms interact exclusively with two actin subunits, each simulation system was simplified to contain one SH3BGRL isoform bound to the two interacting actin subunits. Molecular dynamics simulations were performed using the GROMACS 2024.3 [54] simulation package with the CHARMM36m [55] force field. Each protein complex was placed in a triclinic box with a minimum distance of 1.2 nm between the protein and the box edges to prevent periodic image interactions. The systems were solvated with TIP3P water molecules, and sodium and chloride ions were added to a final concentration of 0.15 M. Additional sodium ions were introduced to neutralize the overall charge of each system. Energy minimization was performed using the steepest descent algorithm to remove steric clashes and unfavourable interactions. The equilibration process was performed in two phases: first, a 100 ps NVT (constant number of particles, volume, and temperature) equilibration was performed to stabilize the temperature at 300 K, followed by a 100 ps NPT (constant number of particles, pressure, and temperature) equilibration using the Parrinello-Rahman barostat to maintain the pressure at 1 bar. For the production simulations, the backbone atoms of both actin subunits were position-restrained with a force constant of 5000 kJ/mol·nm² in all dimensions to maintain the structural integrity of the F-actin pointed end and prevent reorientation during the simulation. Production runs were performed for 500 ns at 300 K using the leap-frog integrator with a 2 fs time step. All bonds involving hydrogen atoms were constrained using the LINCS algorithm. Long-range electrostatic interactions were calculated using the Particle Mesh Ewald method with a real-space cut-off of 1.2 nm, while van der Waals interactions were truncated at 1.2 nm with a force-switch smoothing function applied from 1.0 nm. Coordinates were saved every 10 ps for subsequent analysis.

The root mean square deviation (RMSD) and root mean square fluctuation (RMSF) of the simulation were calculated using the corresponding GROMACS function. The interface analysis was performed using the Python libraries MDAnalysis [56] and ProLIF [57]. The contacts obtained by ProLIF were filtered and transformed into the specific json structure required as Flareplot input using a custom Python script. Visualization of the molecular structure was performed using ChimeraX 1.9 [53].

### Analytical ultracentrifugation

Sedimentation velocity runs were carried out in an Optima AUC analytical ultracentrifuge (Beckman Coulter, USA) using an An-50 Ti rotor at 50,000 rpm and 20 °C. Concentration profiles were measured with the absorption scanning optics at 280 nm or 230 nm using 3 or 12 mm standard double-sector centerpieces filled with 100 μL or 400 μL sample, respectively. Experiments to determine the oligomerization state of SH3BGRL, SH3BGRL-2 and SH3BGRL-3 were performed in a buffer containing 25 mM HEPES pH 7.4, 50 mM NaCl, 0.5 mM TCEP in the indicated concentration range. To test whether SH3BGRL or SH3BGRL-3 are able to interact with G-actin, 10 µM actin supplemented with 15 µM latrunculin B was titrated with the indicated amounts of these proteins in a buffer containing 9 mM Tris pH 7.8, 50 mM KCl, 5 mM MgCl_2_, 0.18 mM CaCl_2_, 45 µM ATP, and 0.18 mM TCEP. After mixing, all samples were allowed to equilibrate for 4 hours at 20 °C prior to sedimentation analysis.

For data analysis a model for diffusion-deconvoluted differential sedimentation coefficient distributions (continuous c(s) distributions) implemented in the program SEDFIT [29] was used. Buffer density, viscosity and partial specific volumes were calculated from the buffer and amino acid composition, respectively, using the program SEDNTERP [58] and were used to correct the experimental s-values to s_20,w_. In the case of mixtures of G-actin and SH3BGRL/-3, experimental s-values are given as s_20,w_ correction is not appropriate due to the different partial specific volumes of the proteins. Figures showing c(s) distributions were obtained using the program GUSSI [59].

Protein concentrations were determined spectrophotometrically using the absorption coefficients at 280 nm as calculated from amino acid composition [60] and are given in monomers throughout the text.

## Data analysis

Data analysis and graph plotting were performed with Origin 2024 (OriginLab Corporation. Massachusetts, USA). Errors are given as standard deviation (SD) based on three independent experiments if not otherwise specified. The significance of the data was evaluated in Origin 2024 using a two-sample t-test (p > 0.05 ≙ ns, p ≤ 0.05 ≙ *, p ≤ 0.01 ≙ **, p ≤ 0.001 ≙ ***, p ≤ 0.0001 ≙ ****).

## Author Contributions

J.N.G. purified proteins and performed biochemical experiments; R.S.H., D.M., and J.N.G. performed AlphaFold predictions and analysis; R.S.H. and D.M. performed molecular dynamics simulations; U.C. performed analytical ultracentrifugation; R.S.H., D.M., U.C. and J.N.G. analyzed the data; R.S.H. and J.N.G designed figures; J.N.G. conceived, coordinated and supervised the study and wrote the manuscript together with R.S.H.

All authors were involved in reviewing and editing the final version of the manuscript.

## Acknowledgments

The authors thank Claudia Thiel and Sandra Witting for excellent technical assistance. We thank Antoine Jégou (Institut Jacques Monod, CNRS, Paris, France) for providing the plasmid encoding mouse SNAP-mDia1(FH1FH2)-His_6_ and Peter Bieling (Max Planck Institute of Molecular Physiology, Dortmund, Germany) for providing the plasmid encoding human His_6_-Tmod3. J.N.G. was supported by the PREPARE program for medical scientists from Hannover Medical School. The Beckman Coulter Optima AUC was co-funded by the DFG and the State of Lower Saxony (INST 192/534-1 FUGG). We would like to thank Dietmar J. Manstein for continuous support and advice and Manuel H. Taft for helpful discussions.

## Materials Availability

Unique reagents generated in this study are available from the lead contact with a completed Materials Transfer Agreement.

---

**Supplementary Figure 1:**
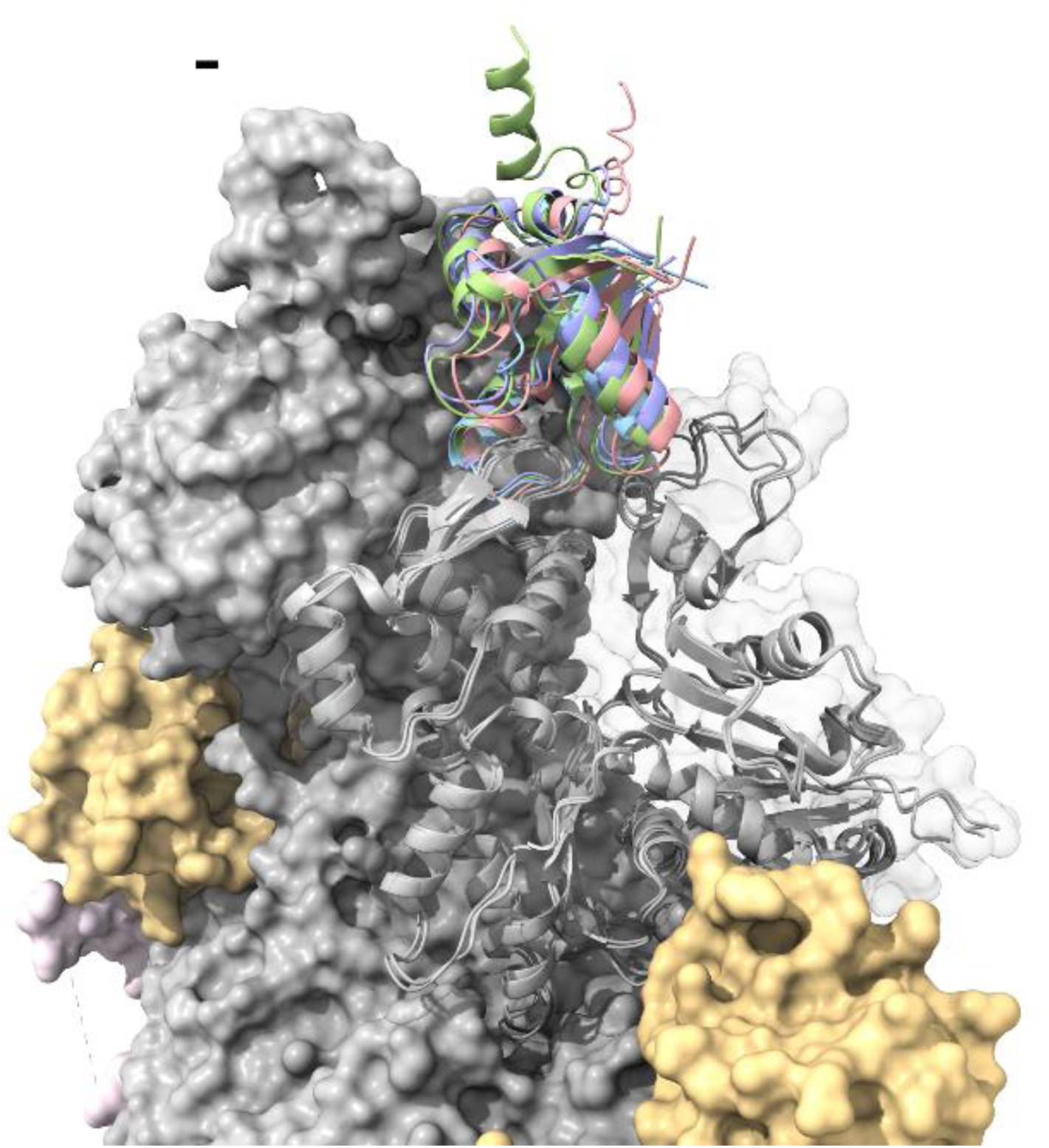
Overlay of the AlphaFold3 generated G-actin–SH3BGRL/-2/-3 models and the published (PDB: 8IAH) structure of the SH3BGRL-2 decorated pointed end in the spectrin–actin complex purified from porcine erythrocytes. Several binding partners of actin were removed from the published structure for the ease of visualization. Grey: actin, yellow: spectrin, green: SH3BGRL, red: SH3BGRL-2, blue: SH3BGRL-3, violet: SH3BGRL-2 in PDB: 8IAH

**Supplementary Figure 2:**
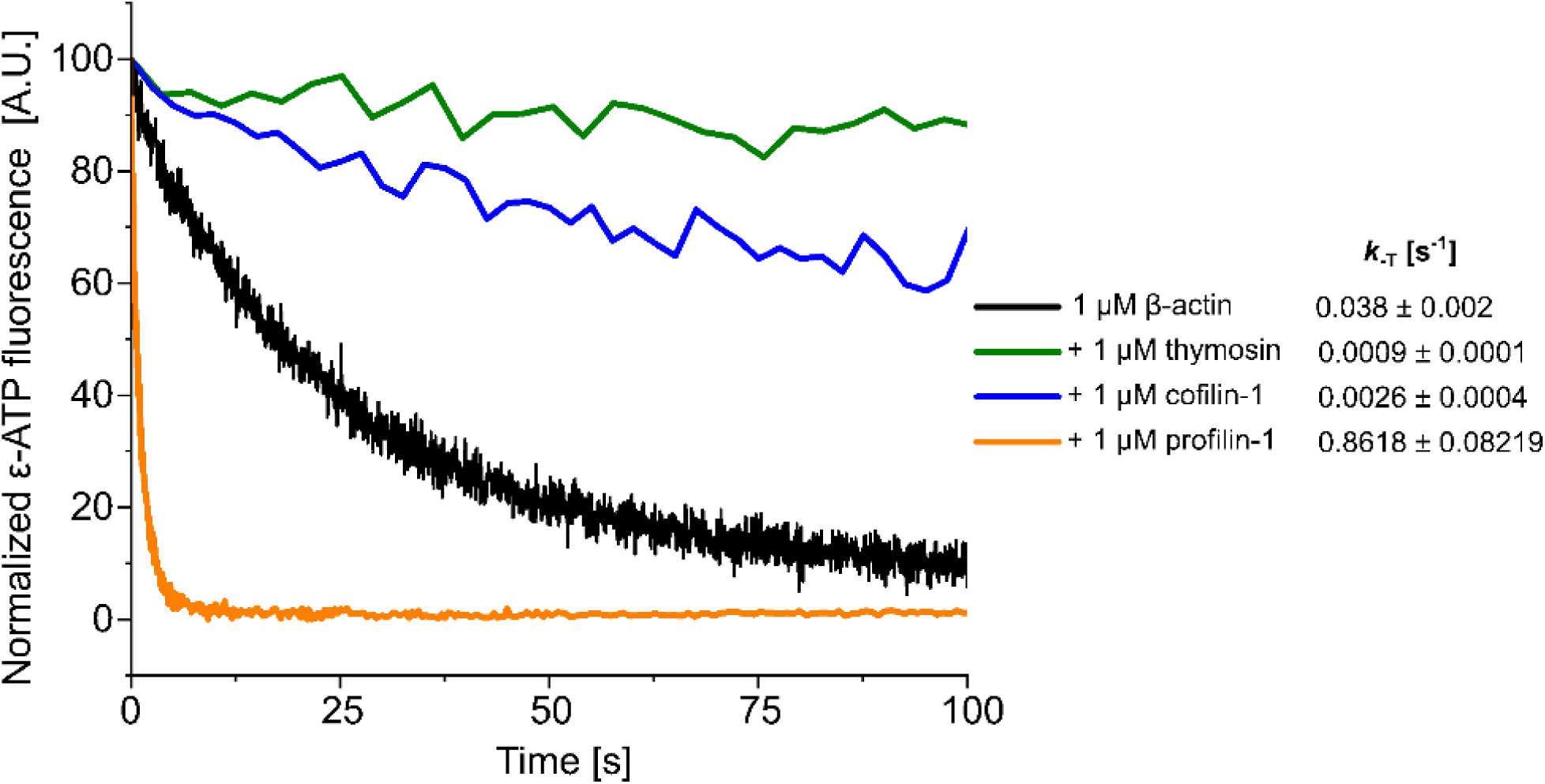
G-actin binding proteins modulate the dissociation of nucleotide from G-actin. The dissociation of fluorescently labeled ATP (ε-ATP) from Mg^2+^-G-actin in the absence and presence of various G-actin binding proteins was followed as described for SH3BGRL-2 (Figure 4A). Shown are representative traces of individual mixing experiments.

**Supplementary Figure 3:**
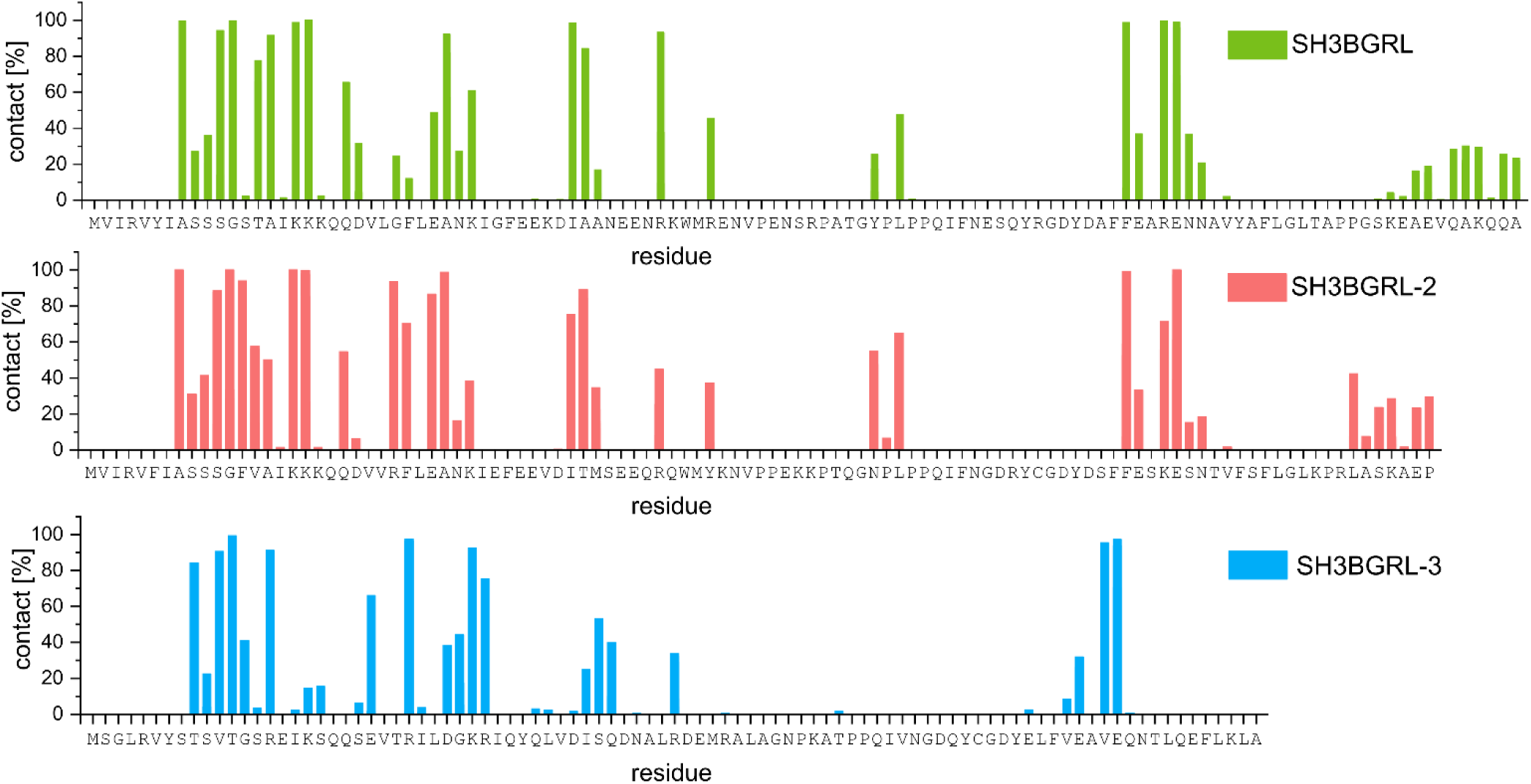
Contact frequency during molecular dynamics simulations. Bar plots of contact frequencies, representing the fraction of the simulation time that the specific residue was in contact with actin.

**Supplementary Figure 4:**
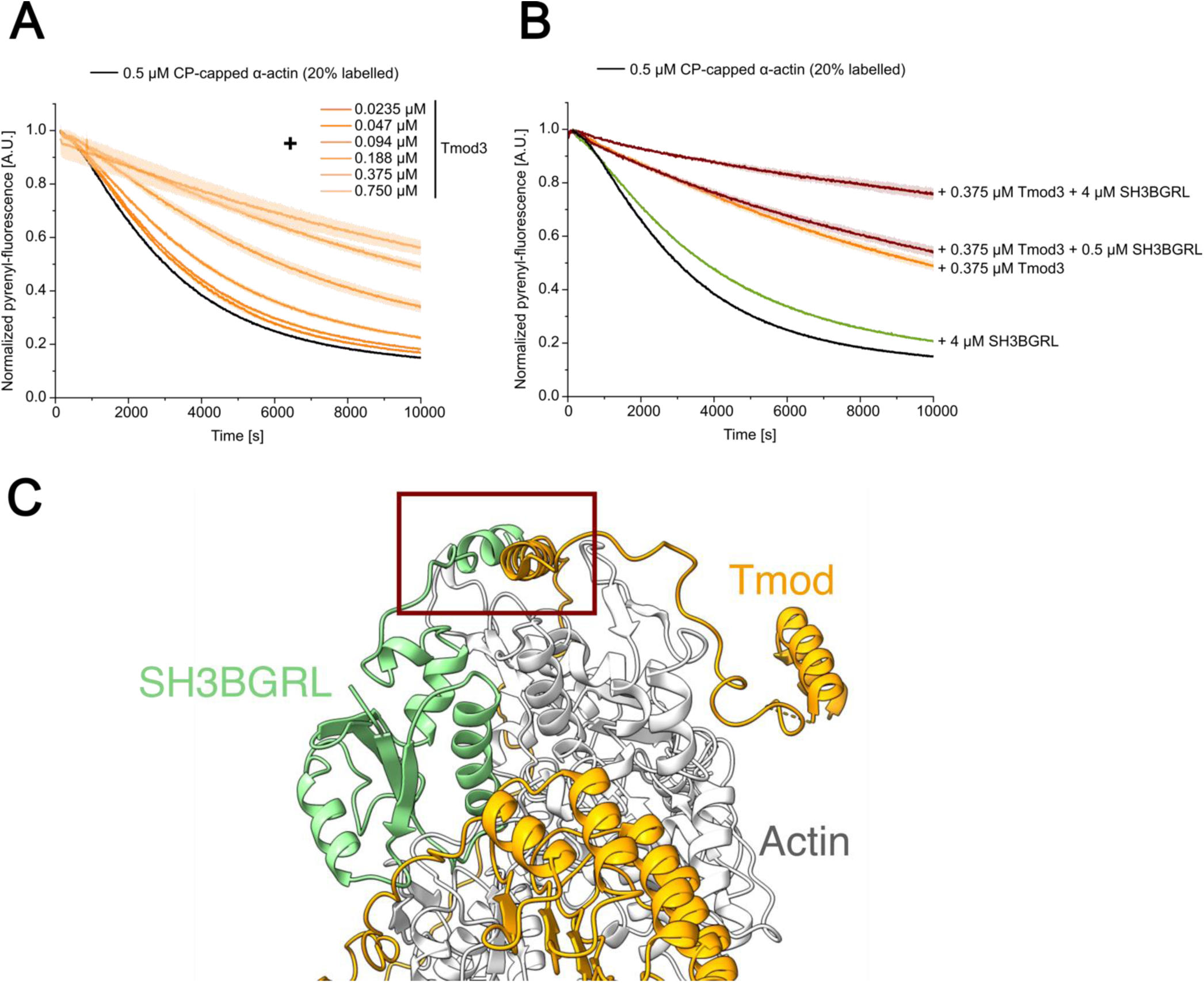
Pyrene-actin based depolymerization experiments of CP-capped filaments in the presence of Tmod3 and/or SH3BGRL. (**A, B**) Traces of dilution-induced depolymerization experiments performed with pyrene-labelled CP-capped actin filaments and (A) increasing concentrations of Tmod3 or (B) mixtures of Tmod3 and SH3BGRL. (**C**) Overlay of the last frame of the SH3BGRL–F-actin MD simulation and the published spectrin–actin complex structure. All other actin binding proteins besides Tmod were removed from the published structure to ease visualization. The possible contact site between the C-terminal region of SH3BGRL and Tmod is indicated. Please note that Tmod1 has been identified in the published structure, but we used the ubiquitously produced isoform Tmod3 for our assays.

**Supplementary Figure 5:**
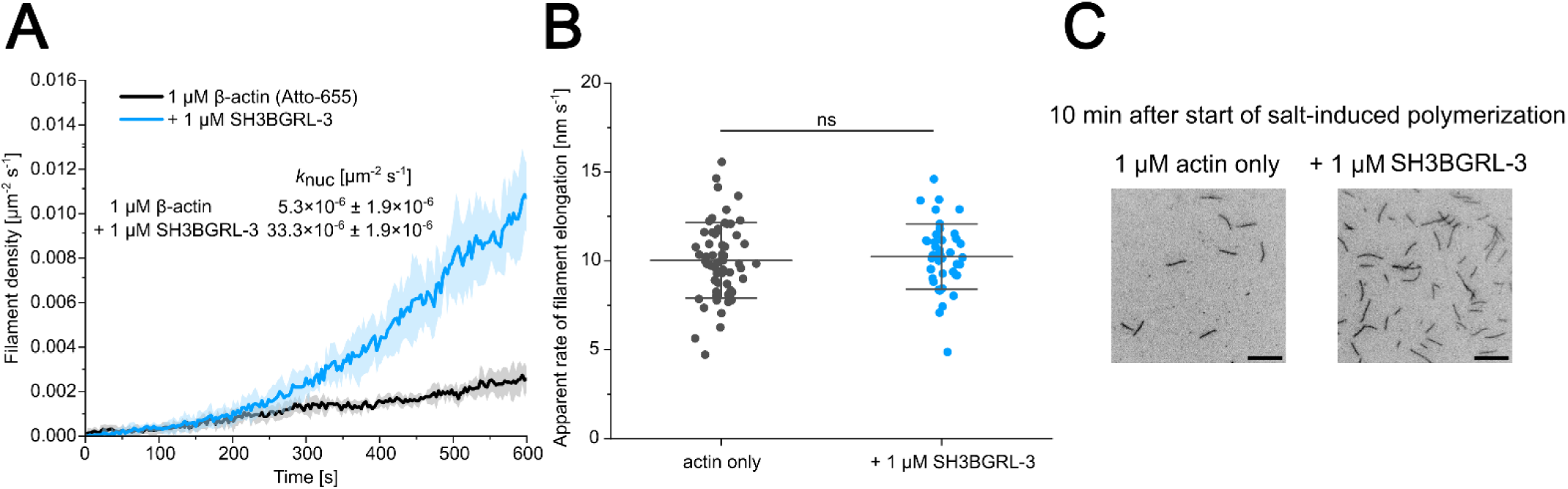
Analysis of the effect of SH3BGRL-3 on nucleation and elongation of recombinant human β-actin using *in vitro* TIRF microscopy (A, **B)** Polymerization of 1 µM fluorescently labeled β-actin (10% Atto-655 labeled) was induced by salt-shift in the absence or presence of 1 µM SH3BGRL-3. The progression of the reaction was tracked by TIRF microscopy. Nucleation efficiency and filament elongation were measured as described in Figure 8. The solid lines and shades in A represent the mean ± SD of at least three individual experiments. Every data point in B represents an individual filament. Data is shown as the mean ± SD. **(C)** Representative micrographs of TIRFM-based polymerization experiments under the indicated conditions 10 min after induction of polymerization. Scale bar corresponds to 10 µm.

**Supplementary Table 1.** Interactors of human SH3BGRL/-2/-3 deposited in the BioGRID database. Only interactors canonically associated with the actin cytoskeleton are shown. Interactors are indicated by their gene name. Database access date: December 2024.

| SH3BGRL | SH3BGRL-2 | SH3BGRL-3 |
| --- | --- | --- |
| ACTB<br>ACTBL2<br>ACTG1<br>ARPC1A<br>ARPC1B<br>ARPC2<br>ARPC3<br>ARPC4<br>ARPC5<br>ARPC5L<br>CAPZA1<br>CAPZA2<br>CAPZB<br>CFL1<br>CORO1B<br>CORO1C<br>CORO2A<br>COTL1<br>CTTN<br>DBN1<br>DSTN<br>GSN<br>LIMA1<br>MYH14<br>MYO1B<br>MYO1C<br>MYO1D<br>MYO1E<br>MYO5A<br>MYO5B<br>MYO5C<br>MYO6<br>PFN1<br>PLS3<br>TMOD3<br>TWF1<br>TWF2<br>WDR1 | ACTB<br>CAPZB<br>GSN<br>MYO1D<br>MYO5A<br>MYO5C<br>TMOD1<br>TMOD2<br>TMOD3 | - |

